# Autofluorescence lifetime imaging classifies human lymphocyte activation and subtype

**DOI:** 10.1101/2023.01.23.525260

**Authors:** Rebecca L. Schmitz, Kelsey E. Tweed, Peter Rehani, Kayvan Samimi, Jeremiah Riendeau, Isabel Jones, Elizabeth M. Maly, Emmanuel Contreras Guzman, Matthew H. Forsberg, Ankita Shahi, Christian M. Capitini, Alex J. Walsh, Melissa C. Skala

## Abstract

New non-destructive tools are needed to reliably assess lymphocyte function for immune profiling and adoptive cell therapy. Optical metabolic imaging (OMI) is a label-free method that measures the autofluorescence intensity and lifetime of metabolic cofactors NAD(P)H and FAD to quantify metabolism at a single-cell level. Here, we investigate whether OMI can resolve metabolic changes between human quiescent versus IL4/CD40 activated B cells and IL12/IL15/IL18 activated memory-like NK cells. We found that quiescent B and NK cells were more oxidized compared to activated cells. Additionally, the NAD(P)H mean fluorescence lifetime decreased and the fraction of unbound NAD(P)H increased in the activated B and NK cells compared to quiescent cells. Machine learning classified B cells and NK cells according to activation state (CD69+) based on OMI parameters with up to 93.4% and 92.6% accuracy, respectively. Leveraging our previously published OMI data from activated and quiescent T cells, we found that the NAD(P)H mean fluorescence lifetime increased in NK cells compared to T cells, and further increased in B cells compared to NK cells. Random forest models based on OMI classified lymphocytes according to subtype (B, NK, T cell) with 97.8% accuracy, and according to activation state (quiescent or activated) and subtype (B, NK, T cell) with 90.0% accuracy. Our results show that autofluorescence lifetime imaging can accurately assess lymphocyte activation and subtype in a label-free, non-destructive manner.

**Teaser:** Label-free optical imaging can assess the metabolic state of lymphocytes on a single-cell level in a touch-free system.

## 1 Introduction

Lymphocytes consist of natural killer (NK) cells, B cells, and T cells, and constitute approximately 20-40% of circulating white blood cells (*1*). T cells cause antigen-specific cytotoxicity and immune-modulating activities after activation, and have been used clinically to treat cancer, viral infections, autoimmune disease, graft-versus-host-disease, and transplant rejection (*2*). NK cells show antigen-independent cytotoxicity and immune-modulating activities after activation, but rely on a balance of activating and inhibitory signals to initiate cytotoxicity (*3*). Similar to T cells, NK cells are emerging in early phase trials as a viable adoptive cell therapy for cancer (*4*), particularly with cytokine induced memory-like NK cells (*5*). B cells, like T cells, are a part of the adaptive immune system, but their primary role is the production of antibodies (*6, 7*). B cells are also antigen-presenting cells that can present peptides to T cells to promote their effector functions (*7, 8*). Subtypes of B cells also secrete cytokines that can either attenuate or suppress the function of surrounding immune cells (*8*). The multiple functions of B cells provide several avenues for leveraging B cells as a platform for cell-based therapies, including antigen-presenting B cells as a cancer immunotherapy and protein production for rare genetic diseases (*6, 9*). Immune profiling of activation of NK, B, and T cells to a stimulus (such as an antigen from a virus or bacterium) can be used to identify the potency of a cell therapy and potentially predict outcome (*10–12*), but current methods are destructive of blood or tissue and require labor-intensive techniques that take hours to days to complete and interpret. New label-free and non-destructive tools are needed to assess lymphocyte activation and subtype in single cells in a more rapid manner.

Single cell measurements capture lymphocyte heterogeneity within a patient, which significantly impacts prognosis (*13–15*). Non-destructive tools enable subsequent analysis and long-term study of cells, while label-free tools enable subsequent expansion and use of cells in patients, for example in adoptive cell therapy (*2, 16*). Current methods to assess lymphocytes include flow cytometry, cytokine release, single-cell RNA sequencing, and cytometry by time of flight (CyTOF). Flow cytometry provides single-cell resolution, but requires labelling with fluorescent antibodies that can be time consuming, may be disruptive to cells, and complicates further use of cells (*17*). Bulk measurements of cytokine release are also popular but do not routinely provide single-cell measurements, and ELISPOT, which provides single-cell cytokine release information also requires cell labeling (*17*). Additionally, cytokine-based techniques cannot provide information about subsets of immune cells that do not secrete cytokines. Finally, single-cell RNA sequencing and CyTOF provide extensive single-cell information, but destroy the sample (*11, 18*).

Optical metabolic imaging (OMI) is an attractive label-free tool to assess the metabolic state of single cells (*19–23*). OMI measures the autofluorescence intensity and lifetime of metabolic cofactors reduced nicotinamide adenine dinucleotide (phosphate) [NAD(P)H] and flavin adenine dinucleotide (FAD) (*24–26*). The fluorescence of NADPH and NADH overlap, and are jointly referred to as NAD(P)H (*25*). Since only the reduced form of NADPH and NADH and the oxidized form of FAD are fluorescent, the fluorescence intensity ratio of NAD(P)H to FAD is defined as the “optical redox ratio”, which provides information about the overall redox state of the cell (*20, 27*). NAD(P)H and FAD each have two distinct fluorescence lifetimes due to their unbound and protein-bound states, so fluorescence lifetime imaging (FLIM) provides insight into changes in unbound and protein-bound pools for each co-enzyme, along with changes in lifetimes due to environmental factors and preferred binding partners (*28–30*). OMI relies on endogenous fluorophores already present in cells, so it is minimally invasive and can provide nondestructive monitoring of cellular metabolism (*28*). Cell segmentation algorithms developed with OMI enable single-cell resolution, which provides insight into metabolic heterogeneity within the population (*31*).

OMI is a promising technique to evaluate lymphocyte activation and subtype because known metabolic shifts occur with activation and between NK, B, and T cells. Unstimulated NK, B, and T cells have low metabolic demands and largely rely on low levels of glycolysis and oxidative phosphorylation to generate ATP (*32–35*). Once activated, extra energy is needed to fuel the effector functions of lymphocytes. In order to fuel rapid proliferation and produce cytokines and other molecules, activated lymphocytes increase use of glucose through aerobic glycolysis and oxidative phosphorylation (*33, 34, 36*). Overnight stimulation with activating cytokines (including IL-2, IL-12, and IL-15) increases rates of glycolysis and oxidative phosphorylation in NK cells (*34, 37, 38*). Similar increases in glycolytic metabolism and oxidative phosphorylation occur with activation in B and T cells (*35, 36*). While these three cell types share a close lineage, the metabolism of NK, B, and T cells are unique. In a study of splenic mouse T and B cells, resting T cells were found to have higher glucose uptake and lactate generation compared to resting B cells, with B cells showing higher mitochondrial mass than T cells (*39*). In T effector cells, fatty acid synthesis is necessary for differentiation and proliferation, but inhibition of this pathway in NK cells does not substantially impact their proliferation (*40, 41*).

Our previous work showed that OMI can classify primary human CD3^+^ and CD3^+^CD8^+^ T cells based on activation status. OMI has also been used to classify subsets of macrophages in monoculture, tumor coculture, and *in vivo* in zebrafish, and to distinguish between categories of blood cells (*i.e*. erythrocytes, monocytes, granulocytes, lymphocytes) (*42–44*). Prior work also shown that NADH autofluorescence intensity increases in activated B cells compared to unstimulated B cells (*45*). These results demonstrate that OMI is promising for lymphocyte profiling, but to our knowledge, no studies to date have built classifiers based on OMI for NK cell activation, B cell activation, or lymphocyte subtype. Given the relevance of NK cells, B cells, and T cells for adoptive cell therapy, this study investigates whether OMI can classify activation in NK cells and B cells, classify lymphocyte subtype (NK, B, T cells), and provide a six-group classifier for activation and lymphocyte subtype. These studies indicate that machine learning classifiers and label-free non-invasive OMI provide high accuracy for single cell classification of activation and lymphocyte subtype from primary human peripheral blood samples.

## 2 Results

### 2.1 OMI resolves metabolic differences between quiescent and activated human B cells

A graphical overview of the experimental design is provided (**Fig. 1A**). Isolated human B cells were activated using anti-CD40 antibody and IL-4 to mimic T cell mediated activation (*46*). After 72 hours of *in vitro* activation, media was collected for cytokine, glucose, and lactate assays, then cells were stained with anti-CD69 to identify activated and quiescent cells in each condition for subsequent OMI. To confirm that our protocol successfully activated B cells, the concentration of IL-6 in the media was measured at 72 hours and found to be significantly increased in the activated compared to the control condition (**Fig. 1B**). Similarly, analysis of glucose and lactate levels at 72 hours show decreased glucose and increased lactate in the media of activated compared to control B cells (**Fig. 1C-D**), confirming known metabolic changes with B cell activation (*35, 36*). Representative images from OMI (**Fig. 1E**) include NAD(P)H mean fluorescence lifetime (τ_m_), FAD τ_m_, optical redox ratio, and CD69 fluorescence images in pseudocolor. Qualitatively, most B cells in the activated condition stain positive for CD69.

**Figure 1.**
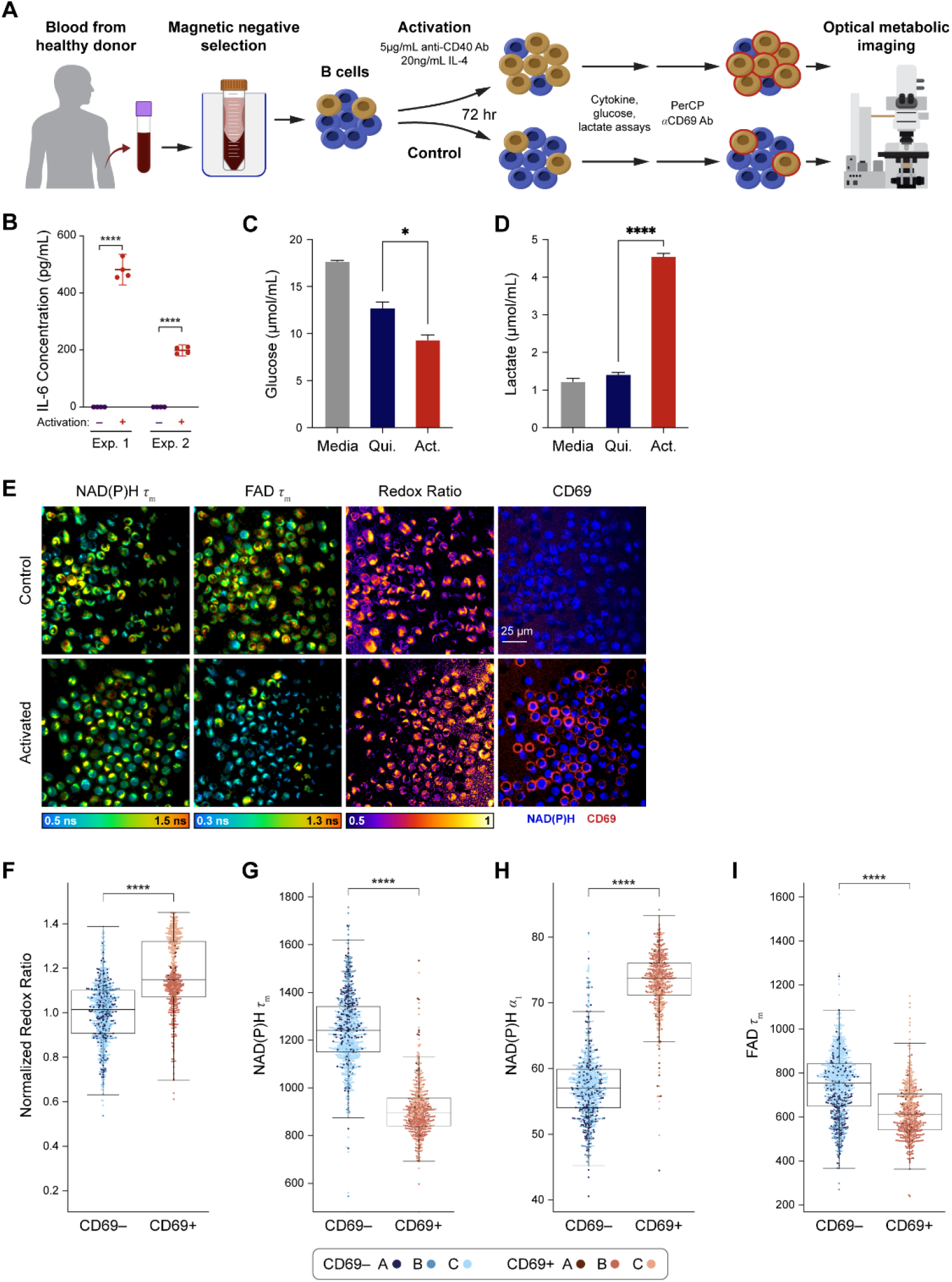
Optical metabolic imaging of primary human B cells activated with IL-4 and anti-CD40. (A) B cells were isolated from human peripheral blood of three different donors and activated for 72 hours with 5 μg/mL anti-CD40 and 20 ng/mL IL-4, or cultured unstimulated. (B) IL-6 concentration was measured in media collected from B cells isolated from two different donors and cultured with or without anti-CD40/IL-4 for 72 hours. The increase in IL-6 concentration in the activated B cell condition is consistent with T-cell dependent B cell activation. **** P < 0.0001, parametric T-test. (C) Samples of media from activated and quiescent B cells were taken before imaging and measured using commercial kits. Glucose in the media of activated B cells was significantly decreased compared to the quiescent cell media. (D) Lactate levels in activated B cell media were significantly higher than lactate levels in the quiescent cell media. (E) Representative images of NAD(P)H τ_m_, FAD τ_m_, redox ratio (NAD(P)H intensity divided by the sum of NAD(P)H and FAD intensity), and anti-CD69 staining in the unstimulated and activated conditions. (F) Redox ratio normalized to the mean of the control group significantly increased in the CD69+ B cells in the IL-4 + anti-CD40 condition compared to CD69-B cells in the unstimulated condition. (G) – (H) NAD(P)H τ_m_ significantly decreased and NAD(P)H α_1_ significantly increased in the CD69+ B cells in the IL-4 + anti-CD40 condition compared to CD69-B cells in the unstimulated condition. (I) A significant decrease in FAD τ_m_ was seen in the CD69+ B cells in the IL-4 + anti-CD40 condition compared to CD69-B cells in the unstimulated condition. In (C) – (D), media samples were diluted 100-fold and 0.5μL was assayed. Assays were performed according to the respective BioVision kit protocols. * P < 0.05, **** P < 0.0001, parametric T-test. In (F) – (I), data are displayed as box-and-whisker plots, representing the median and interquartile range (IQR), with whiskers at 1.5*IQR. Plots are overlaid with data points; each point represents one cell. n = 1210 cells (461 cells in the activated CD69+ condition, 749 cells in the control CD69-condition). **** P < 0.0001, two-tailed unpaired T-test.

The optical redox ratio (NAD(P)H intensity divided by the sum of NAD(P)H and FAD intensity) was elevated in CD69^+^ B cells in the activated condition compared to CD69^−^ B cells in the control condition (**Fig. 1F**). Additionally, NAD(P)H τ_m_ decreased and NAD(P)H α_1_ (the fraction of free, unbound NAD(P)H) increased in CD69^+^ activated B cells compared to the CD69^−^ control cells (**Fig. 1G, 1H**). FAD τ_m_ also decreased in the CD69^+^ activated cells compared to the CD69^−^ control cells (**Fig. 1I**).

When comparing CD69^+^ and CD69^−^ cells within the unstimulated or activated conditions, the CD69^+^ and CD69^−^ B cells in the unstimulated condition did not show any significant differences in OMI parameters. However, in the activated condition, CD69^+^ cells were significantly different compared to CD69^−^ cells for all OMI parameters besides the optical redox ratio (**Supp. Fig. 1**).

### 2.2 Single cell clustering and machine learning models based on OMI separate B cells by activation state

Next, we investigated whether OMI could visualize single cell heterogeneity in B cells and whether machine learning models based on OMI could classify B cell activation state. Unsupervised clustering of 9 OMI parameters from single cells in the CD69^+^ activated condition and CD69^−^ control condition revealed that the CD69^+^ activated cells cluster separately from the CD69^−^ control cells across all three donors (**Fig. 2A**). Uniform manifold approximation and projection (UMAP) was used to visualize the clustering of single B cells based on the same OMI parameters, which similarly revealed distinct clusters of CD69^+^ activated and CD69^−^ control cells (**Fig. 2B**). Additional UMAPs colored by donor, condition (control, stimulated), and CD69 status are provided (**Supp. Fig. 2A-B**).

**Figure 2.**
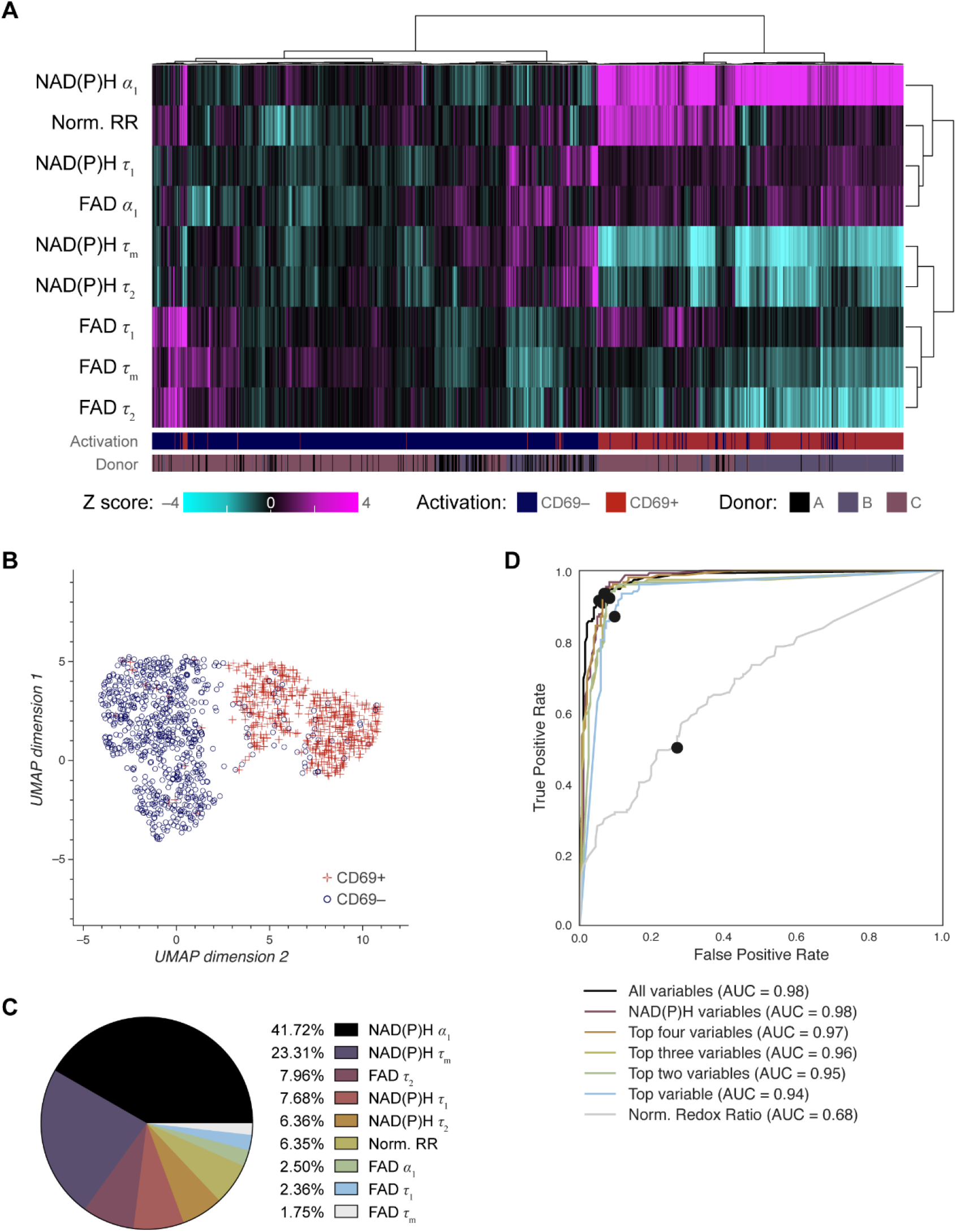
Heterogeneity and classification of activated and quiescent B cells using OMI parameters. (A) Heatmap of single-cell data across all B cell experiments. Hierarchical cell clustering was calculated based on the z-scores (the difference between cell mean and population mean divided by the population standard deviation) of nine OMI variables (NAD(P)H τ_m_, τ_1_, τ_2_, α_1_; FAD τ_m_, τ_1_, τ_2_, α_1_; and control-normalized optical redox ratio). The single-cell clustering demonstrates that using all OMI variables, activated B cells tend to group separately from quiescent B cells regardless of donor. (B) UMAP of nine OMI parameters visualizes separation between clusters of activated (CD69+ in activated condition) and quiescent (CD69- in unstimulated condition) B cells. (C) Pie chart showing the relative weight of the nine OMI variables included in the “all variables” random forest classifier. (D) Receiver operating characteristic (ROC) curve of random forest classifiers trained for classification of quiescent and activated B cells on different combinations of OMI variables, with operating points indicated. “Top variables” classifiers refer to the largest weighted variables in the “all variable” classifier, found in (C). An area under the curve (AUC) of 0.98 is indicative of high performance of the “all variable” classifier and the NAD(P)H variables (NAD(P)H τ_m_, τ_1_, τ_2_, α_1_) classifier. n = 1210 cells (461 cells in the activated CD69+ condition, 749 cells in the control CD69-condition) with a 70/30 split for training and test sets.

Next, a random forest classifier based on OMI parameters for each B cell was trained on 70% of the cells and tested on the remaining 30% of cells to identify activated (CD69+ in activated condition) or quiescent (CD69-in unstimulated condition) B cells. The OMI parameters with the greatest weight in the classification of CD69^+^ and CD69^−^ B cells were NAD(P)H α_1_ (41.72%), NAD(P)H τ_m_ (23.31%), unbound FAD fluorescence lifetime (τ_2_) (7.96%), and unbound NAD(P)H fluorescence lifetime (τ_1_) (7.68%) (**Fig. 2C**). The resulting classifier has an accuracy of 93.4% (**Supp. Fig. 2C, Supp. Table 1**), with a receiver operating characteristic (ROC) area under the curve (AUC) of 0.98 (**Fig. 2D**). Logistic regression and support vector machine (SVM) classification performed similarly to the random forest classifier (**Supp. Fig. 2C-F**). Classification based on the NAD(P)H and FAD phasors at both the laser repetition frequency (80MHz) and its second harmonic (160MHz) predicted B cell activation with 93.9% accuracy (**Supp. Fig. 9A-B**).

### 2.3 OMI resolves metabolic differences between quiescent and activated human NK cells

A graphical overview of the NK experiment is provided (**Fig. 3A**). Isolated primary human NK cells were activated *in vitro* for 24 hours using IL-12, IL-15, and IL-18 as previously described for inducing memory-like NK cells (*5*). After 24 hours of *in vitro* activation, media was collected for cytokine, glucose, and lactate assays, then cells were stained with anti-CD69 to identify activated and quiescent cells in each condition for subsequent OMI. To confirm NK cell activation, the concentration of IFN-γ in the media was measured at 24 hours and found to significantly increase in activated compared to control NK cells (**Fig. 3B**). Similarly, analysis of glucose and lactate levels at 24 hours show decreased glucose and increased lactate in the media of activated compared to control NK cells (**Fig. 3C-D**), confirming known metabolic changes with NK cell activation (*34, 37, 38*). Representative images of NAD(P)H τ_m_, FAD τ_m_, optical redox ratio, and CD69 expression are presented in pseudocolor (**Fig. 3E**). Qualitatively, most NK cells in the activated condition express CD69.

**Figure 3.**
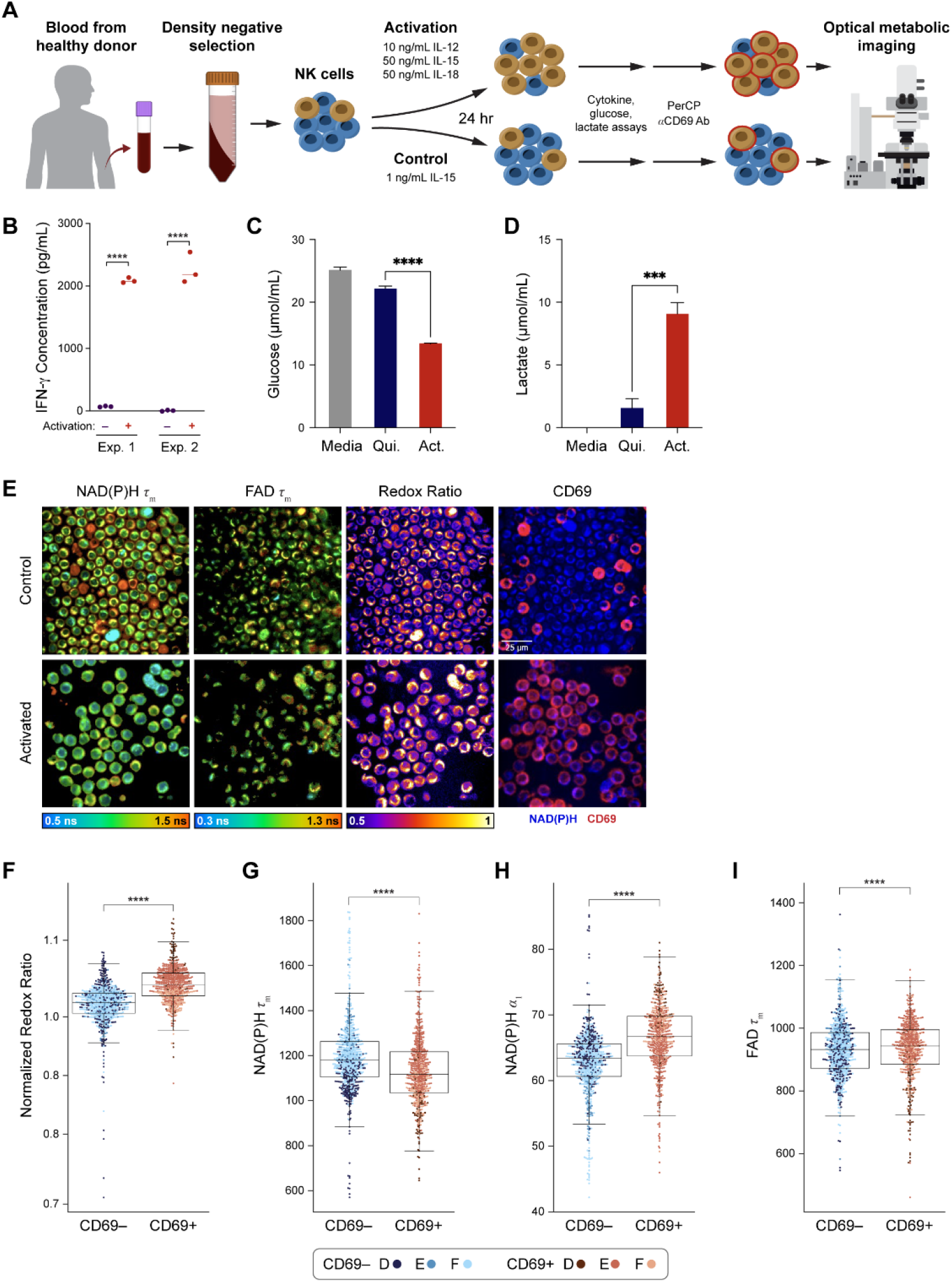
Optical metabolic imaging of primary human NK cells activated with IL-12, IL-15, and IL-18. (A) NK cells were isolated from human peripheral blood of three different donors and activated with 10 ng/mL IL-12, 50 ng/mL IL-15, and 50 ng/mL IL-18 for 24 hours. (B) IFN-y concentration in media collected from NK-cells isolated from two different donors and cultured with or without activating cytokines for 24 hours. The increase of IFN-y in the activated condition is consistent with NK cell activation. **** P < 0.0001, parametric T-test. (C) Samples of media from activated and quiescent NK cells from two different donors were taken before imaging and measured using commercial kits. Glucose in the media of activated NK cells was significantly decreased compared to the quiescent cell media. (D) Lactate levels in activated NK cell media were significantly higher than lactate levels in the quiescent cell media. (E) Representative images of NAD(P)H τ_m_, FAD τ_m_, redox ratio, and anti-CD69 staining in the control and activated conditions. (F) Redox ratio significantly increased in the CD69+ NK cells in the cytokine-activated condition compared to CD69-NK cells in the unstimulated condition. (G) – (I) NAD(P)H τ_m_ significantly decreased, and FAD τ_m_ and NAD(P)H α_1_ significantly increased, in the CD69+ NK cells in the cytokine-activated condition compared to CD69-NK cells in the unstimulated condition. In (C) – (D), media samples were diluted 100-fold and 0.5μL was assayed. Assays were performed according to the respective BioVision kit protocols. *** P < 0.001, **** P < 0.0001, parametric T-test. In (F) – (I), data are displayed as box-and-whisker plots, representing the median and interquartile range (IQR), with whiskers at 1.5*IQR. Plots are overlaid with data points; each point represents one cell. n = 1221 cells (554 cells in the activated CD69+ condition, 667 cells in the control CD69-condition). **** P < 0.0001, two-tailed unpaired T-test.

OMI of NK cells revealed several changes in CD69^+^ NK cells under activating conditions compared to CD69^−^ cells under control conditions. The optical redox ratio significantly increased in CD69^+^ activated NK cells compared to CD69^−^ control NK cells (**Fig. 3F**). NAD(P)H τ_m_ decreased, and NAD(P)H α_1_ and FAD τ_m_ increased in activated NK cells compared to quiescent control NK cells (**Fig. 3G-I**).

OMI parameters were compared across both CD69^+^ and CD69^−^ NK cells under activating and control conditions. Most OMI parameters did not change with CD69 status within activating or control conditions, except the optical redox ratio and NAD(P)H τ_1_ (**Supp. Fig. 3**).

### 2.4 Single cell clustering and machine learning models based on OMI separate NK cells by activation state

Next, we investigated whether OMI could visualize single cell heterogeneity in NK cells and whether machine learning models based on OMI can classify NK cell activation state. Unsupervised clustering of 9 OMI parameters from single cells in the CD69^+^ activated condition and CD69^−^ control condition revealed that NK cells were somewhat heterogeneous, resulting in the emergence of a dominant cluster with several smaller clusters of activated and quiescent cells (**Fig. 4A**). A UMAP was used to visualize the clustering of single NK cells based on the same OMI parameters, which demonstrated a cluster of CD69+ NK cells away from a cluster of a mixed CD69- and CD69+ NK cell population (**Fig. 4B**). Further color-coding by donor reveals that NK cells from all three donors overlap in these clusters (**Supp. Fig. 4B**).

**Figure 4.**
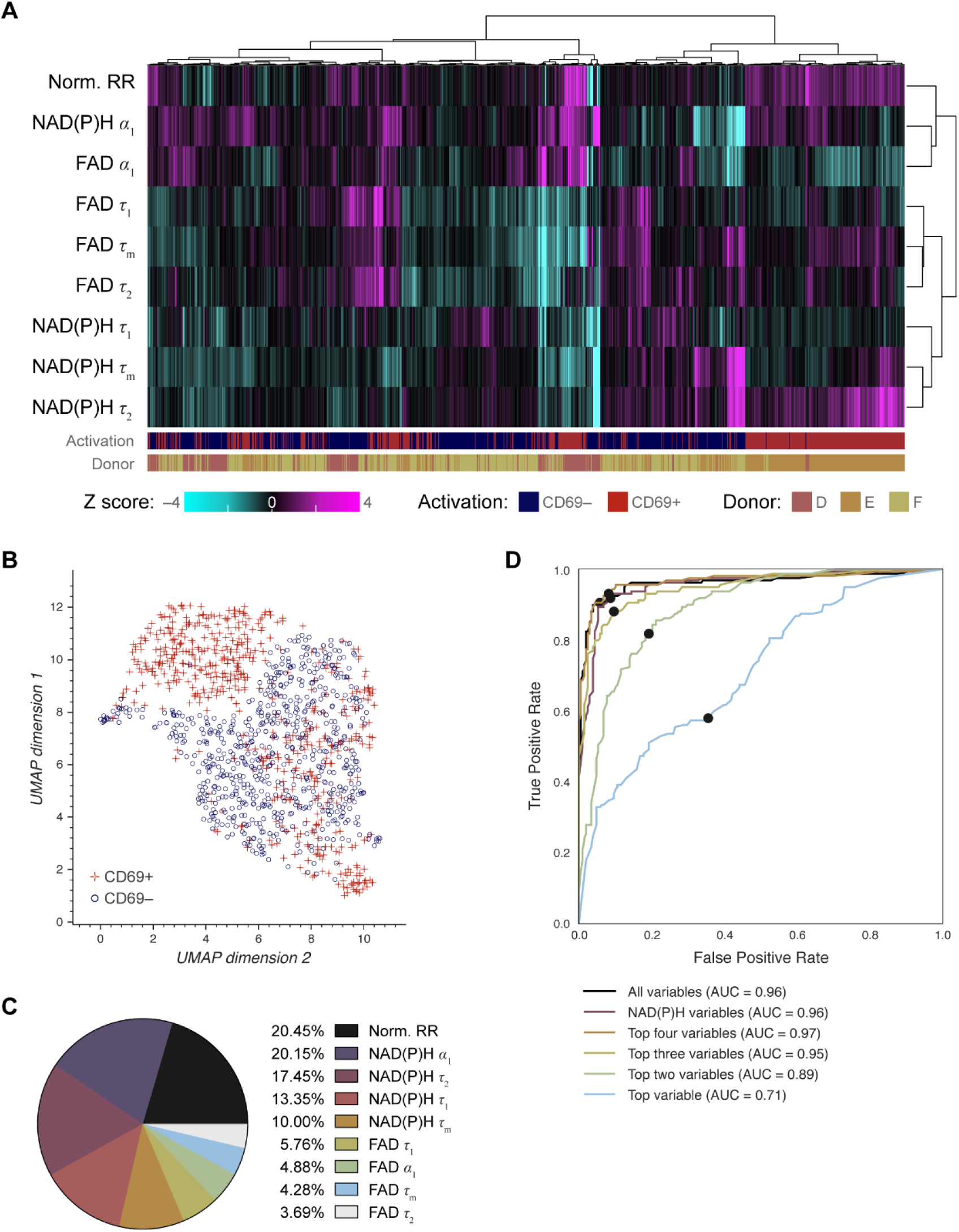
Heterogeneity and classification of activated and quiescent NK cells using OMI parameters. (A) Heatmap of single-cell data across all NK cell experiments reveals heterogeneity within the dataset. Hierarchical cell clustering was calculated on the z-scores (the difference between cell mean and population mean divided by the population standard deviation) of nine OMI variables (NAD(P)H τ_m_, τ_1_, τ_2_, α_1_; FAD τ_m_, τ_1_, τ_2_, α_1_; and control-normalized optical redox ratio). (B) UMAP of nine OMI parameters displays clustering of activated (CD69+ in activated condition) and quiescent (CD69- in unstimulated condition) NK cells. (C) Pie chart showing the relative weight of each of the nine OMI parameters in the “all variable” random forest classifier. (D) ROC curve of random forest classifiers trained for classification of quiescent and activated NK cells based on different combinations of OMI parameters, with operating points indicated. “Top variables” classifiers refer to the largest weighted OMI parameters in the classifier using all variables, displayed in (C). The classifier using the top four OMI parameters performed the best (AUC 0.97), followed by the classifier that used all 9 OMI parameters (AUC 0.96) and the classifier that used only NAD(P)H lifetime variables (NAD(P)H τ_m_, τ_1_, τ_2_, α_1_) (AUC 0.96). n = 1221 cells (554 cells in the activated CD69+ condition, 667 cells in the control CD69-condition) with a 70/30 split for training and test sets.

A random forest classifier based on single-cell OMI parameters was trained and tested on 70% and 30%, respectively, of the NK cells to identify activated (CD69+ in activating conditions) or quiescent (CD69- in unstimulated conditions) states. The highest weighted OMI parameters were the control-normalized optical redox ratio (20.45%), NAD(P)H α_1_ (20.15%), protein-bound NAD(P)H fluorescence lifetime (τ_2_) (17.45%), and unbound NAD(P)H fluorescence lifetime (τ_1_) (13.35%) (**Fig. 4C**). The resulting classifier had an accuracy of 92.6% (**Supp. Fig. 4C, Supp. Table 1**), and the AUC of the ROC curve was 0.96 (**Fig. 4D**). Logistic regression and SVM classification had a slightly lower performance than random forest classification, with AUC of the ROC curves of 0.95 and 0.94, respectively (**Supp. Fig. 4C-F**). Classification based on the NAD(P)H and FAD phasors at both the laser repetition frequency (80MHz) and its second harmonic (160MHz) predicted NK cell activation with 89.2% accuracy (**Supp. Fig. 9C-D**).

### 2.5 OMI quantifies lymphocyte heterogeneity and classifies lymphocyte subtype and activation state

We then investigated whether OMI parameters could distinguish activation and/or lymphocyte subtype across a dataset containing multiple subtypes of lymphocytes. We combined the NK cell and B cell data with our previously published T cell data (quiescent and activated for 48 h with CD2/3/28) (*47*) and plotted several key OMI parameters, including the control-normalized optical redox ratio, NAD(P)H τ_m_, and NAD(P)H α_1_ (**Fig. 5A**). Across all three lymphocyte subtypes, these variables exhibited similar changes with activation: NAD(P)H τ_m_ decreased with activation, while NAD(P)H α_1_ and the optical redox ratio increased with activation. These changes with activation were statistically significant in all cases. In addition to activation-associated shifts in OMI parameters, there were also statistically significant differences between quiescent T, B, and NK cells (**Fig. 5A**).

**Figure 5.**
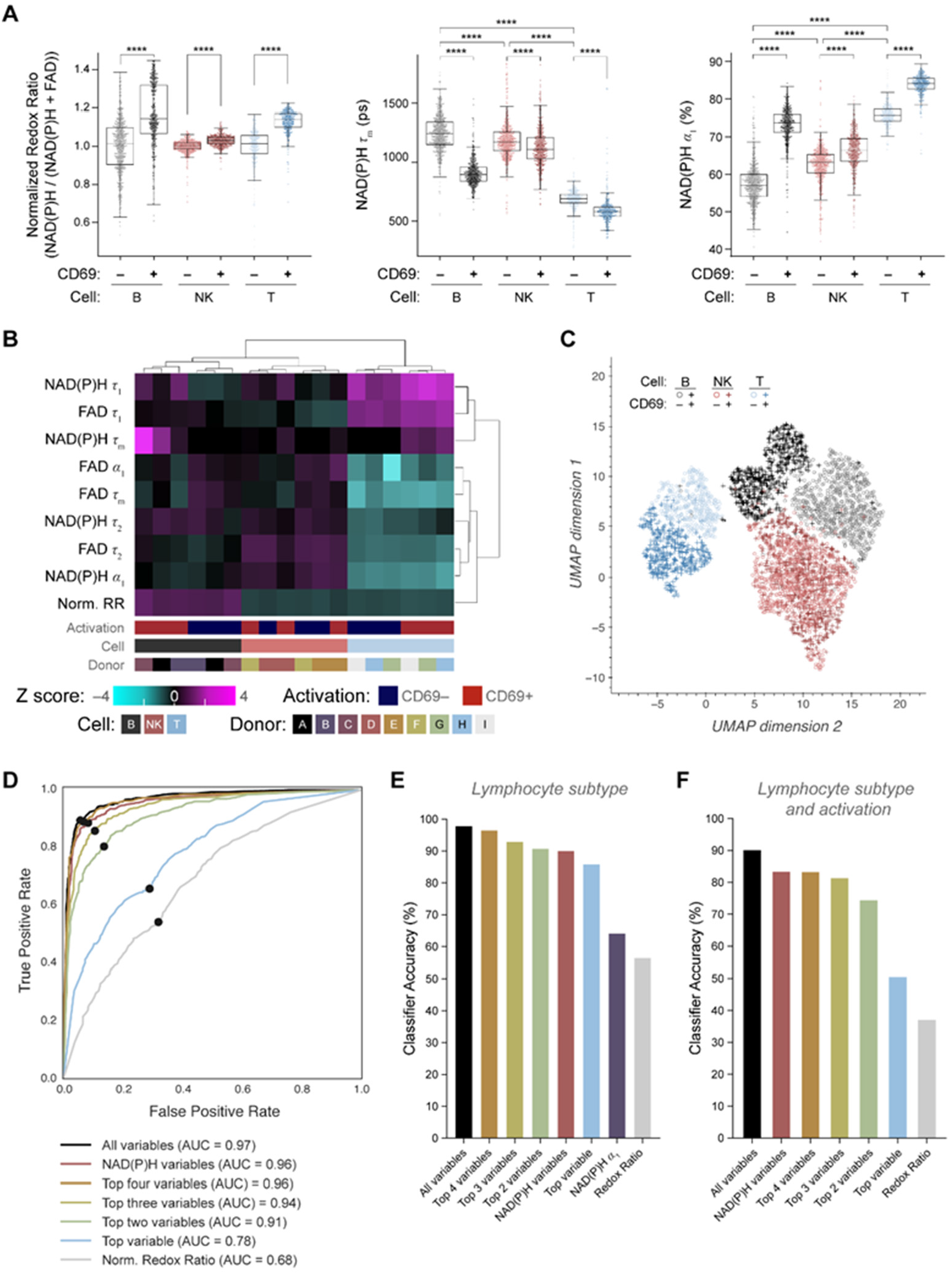
Classification of lymphocyte activation status based on OMI parameters collected on B cells, NK cells, and T cells. Data from activated and quiescent T cells, B cells, and NK cells was used to evaluate OMI measurements across lymphocytes. T cell data from our prior work (*47*) where T cells were activated with CD2/3/28 for 48h and imaged with OMI. (A) Box-and-whisker plots of key OMI variables (control-normalized optical redox ratio, NAD(P)H τ_m_, and NAD(P)H α_1_) display consistent changes with activation across T cells, B cells, and NK cells. Additional changes were noted between quiescent (CD69-control) cells in each of the three lymphocyte subtypes (comparisons between quiescent groups were interpreted as not meaningful for the optical redox ratio, due to normalization). (B) Heatmap displaying hierarchical clustering of groups of activated or quiescent cells by lymphocyte subtype, donor, and activation status, calculated from the z-scores (the difference between experimental group mean and the mean of all cells divided by the standard deviation of all cells) of nine OMI variables. (C) UMAP of single-cell OMI data displays distinct clusters of lymphocytes based on lymphocyte subtype and activation status. (D) ROC curves of random forest classifiers trained to identify activated cells across all three lymphocyte subtypes, with operating points indicated. The highest weighted OMI parameters were used in the “top variables” classifiers; these weights are in Supp. Fig. 5C. (E) Accuracy of different random forest classifiers trained to identify lymphocyte subtype (one vs one approach). Variable weights for “top variables” are in Supp. Fig. 6B. (F) Accuracy of random forest classifiers trained to identify lymphocyte subtype and activation across all three lymphocyte subtypes (one vs. one approach) using different OMI parameters. Variable weights are in Supp. Fig. 8B. n = 3127 cells (749 CD69-control B cells, 461 CD69+ activated B cells, 667 CD69-control NK cells, 554 CD69+ activated NK cells, 331 CD69-control T cells, 365 CD69+ activated T cells) with a 50/50 split for training and test sets. **** P < 0.0001, Kruskal-Wallis with post-hoc comparisons. ns = not significant.

We next used the combined data set of T, B, and NK cells to visualize heterogeneity between each group. Unsupervised clustering was performed across 9 OMI parameters using averages from CD69^+^ activated and CD69^−^ control lymphocytes across activation state (CD69+, CD69-), donor, and lymphocyte subtype (B, NK, T cell) (**Fig. 5B**). This revealed distinct clusters of CD69- and CD69+ T cell and B cell groups. Within the NK cells, clustering was mixed across CD69- and CD69+ status and donors. A UMAP was also used to further visualize clustering, with T cells forming a distinct cluster from B cells and NK cells (**Fig. 5C**).

A UMAP reveals that CD69+ lymphocytes clustered somewhat separately from CD69-lymphocytes (**Supp. Fig. 5A**). Therefore, we investigated whether machine learning models could classify activation within the combined lymphocyte data. First, random forest classification was used to identify whether cells were activated (CD69+) or quiescent (CD69-). Using all 9 OMI parameters, an ROC AUC of 0.97 (**Fig. 5D**) and accuracy of 92.2% (**Supp. Fig. 5B, Supp. Table 1**) was achieved. The top feature weights were NAD(P)H α_1_ (27.10%), NAD(P)H τ_1_ (14.61%), control-normalized optical redox ratio (14.35%), and NAD(P)H τ_2_ (12.60%) (**Supp. Fig. 5C**). Classification based on NAD(P)H lifetime variables (τ_m_, τ_1_, τ_2_, α_1_) alone also had a high ROC AUC of 0.96 (**Fig. 5D**) and performance (accuracy = 90.3%) (**Supp. Fig. 5B, Supp. Table 1**). Logistic regression and support vector machine classification performance were somewhat diminished from the random forest classification performance, with accuracies of 84.8% and 81.5% respectively (**Supp. Fig. 5D-G**).

Similarly, a UMAP reveals that T cells, B cells, and NK cells clustered separately (**Supp. Fig. 6A**). Therefore, three-class random forest classification of lymphocyte subtype was then performed (one vs. one approach) using different combinations of OMI parameters. Feature weights are provided (**Supp. Fig. 6B**). Classification with all nine OMI parameters performed the best (accuracy = 97.8%) (**Fig. 5E**, **Supp. Fig. 6C, Supp. Table 1**). However, other random forest classifiers with fewer parameters also demonstrated strong performance. The top four parameters (FAD τ_1_, FAD τ_m_, NAD(P)H τ_m_, and FAD α_1_) had an accuracy of 96.4%, while NAD(P)H lifetime variables (τ_m_, τ_1_, τ_2_, α_1_) had an accuracy of 89.9% (**Fig. 5E, Supp. Table 1**). Classification using both NAD(P)H and FAD 80MHz and 160 MHz phasors classified cells as either B or NK cells with 99.9% accuracy (**Supp. Fig. 9E-F**), while the NAD(P)H 80MHz and 160MHz phasor classified cells as B, NK, or T cells with 96.2% accuracy (**Supp. Fig. 10A-B**). Classification was also tested on a subset of the combined lymphocyte data that contained only quiescent (CD69-) cells (UMAP, **Supp. Fig. 7A**), and had similarly high performance with all nine OMI parameters (accuracy = 98.4%) (**Supp. Fig. 7B-D, Supp. Table 1**).

Finally, a UMAP shows that quiescent and activated T cells, B cells, and NK cells clustered separately (**Supp. Fig. 8A**). Therefore, random forest classification was used to classify both lymphocyte subtype and activation simultaneously. A six-class classification was performed (one vs. one approach). Feature weights are provided (**Supp. Fig. 8B**). Again, the classifier with all 9 OMI parameters had the highest accuracy (accuracy = 90.0%) (**Fig. 5F, Supp. Table 1**), and misclassification was highest between quiescent vs. activated cells within a lymphocyte subtype, with lymphocyte subtype usually identified correctly (**Supp. Fig. 8C**). Other classifiers also performed well, including the top four parameters (NAD(P)H α_1_, FAD τ_1_, NAD(P)H τ_1_, FAD τ_m_) with an accuracy of 83.2%, and NAD(P)H lifetime variables (τ_m_, τ_1_, τ_2_, α_1_) with an accuracy of 83.3% (**Fig. 5F, Supp. Table 1**). Classification using NAD(P)H 80MHz and 160MHz phasor classified both lymphocyte subtype and activation simultaneously with an accuracy of 88.5% (**Supp. Fig. 10C**). A summary of the accuracies of all random forest classifiers is provided (**Supp. Table 1**).

**Table 1.** Isolation and activation conditions for each lymphocyte subtype.

| Lymphocyte subtype | Negative Isolation Kit | Control Medium | Activation Medium | Activation time |
| --- | --- | --- | --- | --- |
| B cell | EasySep Human Naïve B Cell Isolation Kit (StemCell Technologies) | RPMI + 5% FBS + 1% penicillin/streptomycin | Control medium + 5 µg/mL anti-CD40 antibody + 20 ng/mL IL-4 | 72 hours |
| NK Cell | MACS Human NK Cell Isolation Kit (Miltenyi Biotec) | TheraPeak X-VIVO-10 medium (Lonza) + 10% human serum AB + 1 ng/mL IL-15 | Control medium + 10 ng/mL IL-12 + 50 ng/mL IL-15 + 50 ng/mL IL-18 | 24 hours |

## 3 Discussion

Several areas of research and clinical care rely on lymphocyte assessments, but these efforts would benefit from a non-destructive, single-cell, touch-free technology to assess lymphocyte subtype and activation state, which would reduce the cost and time for analysis of heterogeneity within a patient while enabling subsequent study and use of unperturbed cells. In this report, we have demonstrated that OMI is sensitive to metabolic changes that occur with activation in primary human B cells and NK cells. Additionally, machine learning models trained on single-cell OMI parameters can reliably classify quiescent cells in both CD40/IL4 activated B cells and IL12/IL15/IL18 activated memory-like NK cells, as well as distinguish lymphocyte subtypes (NK, B, T cells) and activation within a combined dataset of NK, B, and T cells.

Interestingly, both B cells and NK cells had similar changes in OMI parameters under activation compared to quiescent cells. The optical redox ratio increased, NAD(P)H τ_m_ decreased, and NAD(P)H α_1_ increased in the activated cells (**Figs. 1, 3**). These trends are consistent with our prior work with primary human T cell activation (*47*). The similarity of changes between the B cells, NK cells, and T cells is likely related to similar shifts in metabolism when all three lymphocyte subtypes are activated. All three types of lymphocytes upregulate oxidative phosphorylation and aerobic glycolysis when activated to fuel rapid growth and proliferation (*34, 35, 48*).

Previous studies have demonstrated that alterations to glycolysis significantly affect OMI measurements (*30, 47, 49, 50*). Specifically, inhibition of glycolysis with 2DG selectively reduced the optical redox ratio in activated T cells, indicating that glycolysis is a key regulator of the optical redox ratio in these cells (*47*). Further, the optical redox ratio has a positive correlation (Pearson’s R = 0.89) with the glycolytic index of breast cancer cells (*49*). In this study, measurements of glucose and lactate levels in media from control and activated B and NK cells revealed that the glucose concentration significantly decreased and the lactate concentration significantly increased with activation (**Figs. 1, 3**). This observation is consistent with upregulation of aerobic glycolysis noted in the literature, as well as our observed increase in the optical redox ratio with activation (*33–38, 40*).

The single-cell resolution of OMI makes it a powerful tool for studying and characterizing population heterogeneity. Here, we characterized heterogeneity within activated and quiescent B cell and NK cell populations. Our results demonstrate that within a population of peripheral human B cells or NK cells exposed to the same conditions, cell outcomes may vary. Examination of CD69 expression revealed that there was a mixture of CD69+ and CD69-cells within each group despite exposure to the same media conditions. We chose to focus the analysis on cells that we could confirm to be quiescent (*i.e*., CD69-cells in the quiescent condition) and cells that we could confirm to be activated (*i.e*., CD69+ cells in the activated condition) to better characterize the ability of OMI to assess these cells without complications that could arise from differing cell states. However, OMI did capture differences in NAD(P)H and FAD autofluorescence between activated and quiescent cells within each condition (**Supp. Figs. 1, 3**). Overall, this study demonstrates the sensitivity of OMI to metabolic differences within a heterogenous cell population.

Classifiers based on single-cell OMI accurately identified activation state with up to 93.4% accuracy for B cells and up to 92.6% accuracy for NK cells (**Supp. Table 1**). In addition to classifying activation within a lymphocyte subtype, OMI also classified activation with high accuracy from a combined dataset of T cells, B cells, and NK cells (accuracy = 92.2%, 9 OMI parameters) (**Supp. Fig. 5B, 5E, Supp. Table 1**). We also found that all 9 OMI parameters accurately classified lymphocyte subtype (accuracy of 97.8%, **Fig. 5E, Supp. Fig. 6C, Supp. Table 1**), which reflects the distinct fluorescence lifetimes of NAD(P)H and FAD for different lymphocyte subtypes (**Fig. 5A**). Indeed, a previous study has shown that NAD(P)H and FAD autofluorescence differs between different types of murine white blood cells (including B cells and T cell subtypes) (*45*). OMI parameters also distinguished between lymphocyte subtypes when only quiescent (CD69-control) cells were used (accuracy of 98.4%, **Supp. Fig. 7B, Supp. Table 1**). Differences in NAD(P)H and FAD fluorescence lifetimes between quiescent lymphocytes may be explained by differences in resting cell metabolism between T cells, B cells, and NK cells, which has been observed in previous human and murine studies (*39–41*). Surprisingly, even a six-group classifier of both activation state and lymphocyte subtype achieved high accuracy (90.0%, **Fig. 5F, Supp. Fig. 8C, Supp. Table 1**) which reflects subtle changes in metabolic state for these six classes (*33–41*).

Although a complete set of NAD(P)H and FAD intensities and lifetimes were collected in this study, all 9 OMI parameters may not be necessary for accurate classification. NAD(P)H lifetime variables alone accurately classified activation within B cells (92.6% accuracy), activation within NK cells (91.6%), activation within the combined dataset of B cells, NK cells, and T cells (90.3% accuracy), lymphocyte subtype (89.9% accuracy), and a six-class classifier of both activation and lymphocyte subtype (83.3% accuracy) (**Supp. Table 1**), while the NAD(P)H phasor alone classified B cell activation and NK cell activation with accuracies of 93.9% and 88.1% respectively (**Supp. Fig. 9A-D**), lymphocyte subtype with 96.2% accuracy (**Supp. Fig. 10A-B**), and both activation and lymphocyte subtype with an accuracy of 88.5% (**Supp. Fig. 10C**). This indicates that simplified hardware with only NAD(P)H excitation and emission capabilities would perform as accurately as a two-color NAD(P)H and FAD imaging system, which is an important consideration in the design of simplified hardware for use in clinical labs.

Overall, these studies indicate that OMI can robustly classify lymphocyte activation status and discriminate B cells, NK cells, and T cells. This label-free single-cell imaging and classification approach could have significant implications in cell manufacturing, where in-line technologies are needed to maintain high potency and safety, or in clinical labs where immune cell profiling is needed to inform treatment decisions. The non-invasive nature of this approach also enables time-course studies of lymphocyte function and *in vivo* studies of lymphocytes in a native context.

## 4 Materials and Methods

### 4.1 Isolation of primary human lymphocytes

Primary human lymphocytes were isolated from peripheral blood obtained from healthy adult donors under approval by the UW-Madison Institutional Review Board. After obtaining informed consent from the donors, 10 to 50 mL whole blood was drawn using a sterile syringe with heparin. B cells and NK cells were then isolated from donor whole blood using negative isolation kits.

For NK cells, blood was mixed in a 1:1 ratio with 1X PBS. The peripheral blood:PBS mixture was then overlayed dropwise onto 15 mL of Lymphoprep (STEMCELL Technologies) in 50 mL conical tubes and centrifuged at 400 xg for 30 minutes at 20°C with slow acceleration and no breaks. After centrifugation, the PBMC layer was moved to a fresh 50 mL conical tube using a 10 mL serological pipette, with 35 mL 1X PBS added to each tube. Cells were centrifuged at 400 xg for 10 min at 20°C with normal acceleration and breaks. After centrifugation supernatant was aspirated and cell pellets were resuspended in 5 mL of ACK lysing buffer (Quality Biological) and let sit at room temperature for 5 minutes. ACK lysis reaction was then quenched with 30 mL of 1X PBS per 50 mL tube. Cells were centrifuged at 400 xg for 10 min at 20°C with normal acceleration and breaks. Supernatant was aspirated, with the pellets being combined in 40 mL 1X PBS and passed through a 70 μm filter. PBMCs were counted on the Z1 Particle Counter (Beckman Coulter) by adding 10 μL of the PBMC solution to 10 mL of Isoton II diluent (Beckman Coulter) in a 20 mL cuvette. PBMCs were then labelled with the human NK Cell Isolation Kit (Miltenyi Biotec), with subsequent NK cell isolation using the “depletes” program on an autoMACS Pro Separator and collecting the negative fraction. The isolated cells were then transferred to a cell culture flask or well plate for culture.

For the B cell isolation (EasySep, STEMCELL Technologies), peripheral blood mononuclear cells (PBMCs) were first isolated by diluting the blood with an equal volume of DPBS + 2% FBS, then centrifuging at 1200 xg for 10 minutes in SepMate tubes containing a layer of Lymphoprep. The isolated PBMCs were then washed with DPBS + 2% FBS and centrifuged at 100 xg for 10 minutes. The resulting sample was resuspended to a concentration of 50 million cells/mL in EasySep Buffer (STEMCELL Technologies). 50 μL/mL isolation cocktail and 50 μL/mL cocktail enhancer were added to the sample, according to the EasySep protocol. 50 μL/mL RapidSpheres solution was then added, and the sample was transferred to a magnet for 3 minutes. The enriched B cells were poured into a new tube and the sample was again placed into a magnet for 1 minute. The enriched B cell population was then washed with culture medium and transferred to a cell culture flask or well plate for culture.

### 4.2 Lymphocyte activation and culture

NK cells were cultured in TheraPeak X-VIVO-10 medium (Lonza) supplemented with 10% human serum AB (Sigma Aldrich) and 1ng/mL IL-15 (Biolegend). B cells were cultured in RPMI containing 5% fetal bovine serum and 1% penicillin-streptomycin. Following isolation, each cell population was divided into two groups: a control population cultured in normal medium, and an activated population cultured in control medium supplemented with additional components. NK cell activating medium was supplemented with 10 ng/mL IL-12 (Invivogen), 50 ng/mL IL-15, and 50 ng/mL IL-18 (Biolegend) (*51, 52*). B cell activating medium was supplemented with 5 ug/mL anti-CD40 antibody (R&D systems) and 20 ng/mL IL-4 (R&D Systems) (*6, 53*).

The cells were cultured separately in activating or control medium for a number of hours depending on the lymphocyte subtype; B cells were activated for 72 hours, and NK cells for 24 hours (*34, 45, 53*). Cells were seeded at a density of 1 million cells/mL medium. At the end of the activation time, a sample of growth medium from each group was taken for cytokine analysis. A summary of the isolation and activation conditions used is provided in **Table 1**.

### 4.3 Staining with PerCP conjugated anti-CD69 antibody

At the end of the activation period, cells were stained with anti-CD69 PerCP-conjugated antibody to distinguish activated and quiescent cells within each population (*38, 53*). The cells were centrifuged at 300 xg for 8 minutes, then resuspended to a concentration of 10 million cells/mL medium. 5μL/million cells PerCP-conjugated anti human CD69 antibody (Biolegend) was added to the sample. The cells were then incubated for 30 minutes at room temperature. Following incubation, the cells were washed twice with media and centrifuged at 300 xg for 8 minutes to remove excess antibody from the sample.

### 4.4 Fluorescence lifetime imaging of lymphocytes

For imaging, B cells and NK cells were plated 1 hour before imaging on poly-D-lysine coated glass-bottomed dishes (MatTek) at a seeding density of 200,000 cells in 50 μL media. The cells were imaged with a custom-built multiphoton fluorescence microscope (Ultima, Bruker) using a 100x (NA = 1.45) oil immersion objective and time-correlated single photon counting electronics (SPC-150, Becker & Hickl GbH, Berlin, Germany). The laser (Insight DS+, Spectra-Physics Inc., Santa Clara, CA, USA) was tuned to 750 nm for NAD(P)H excitation, 890 nm for FAD excitation, and 980 nm or 1040 nm excitation for PerCP. Fluorescence emission was detected using a H7422PA-40 GaAsP photomultiplier tube (Hamamatsu Corporation, Bridgewater, NK, USA) and isolated using a 440/80 bandpass filter for NAD(P)H, 550/50 (NK cells) or 550/100 (B cells) bandpass filter for FAD, and 690/50 bandpass filter for PerCP. In the B cell experiments, the laser power at the sample was 1.5 mW – 2.0 mW for NAD(P)H, 3.0 mW – 4.0 mW for FAD, and 3.0 mW for PerCP. In the NK-cell experiments, the laser power at the sample was 2.0 mW for NAD(P)H, 5.0 mW for FAD, and 3.5 mW for PerCP. The laser power was maintained at a consistent value within each experiment.

300 μm × 300 μm fluorescence lifetime images (256×256 pixels) were collected consecutively for NAD(P)H and FAD in the same field of view, with a pixel dwell time of 4.8 μs and an integration time of 60s. An instrument response function was collected during imaging from the second harmonic generation of a urea crystal, and photon count rates were maintained around 1×10^5^ photons. An intensity image of PerCP fluorescence was collected for the same field of view. Images were collected from three to six fields of view for each sample.

### 4.5 Image analysis

Fluorescence lifetimes were extracted through analysis of the fluorescence decay at each pixel in SPCImage (Becker & Hickl). To provide more robust calculations of the fluorescence lifetimes, a threshold was used to exclude background pixels with a low intensity, and images were binned up to a bin factor of 3 to reach a peak of at least 100 photons in the decay. Both NAD(P)H and FAD can exist in a quenched and an unquenched configuration with distinct lifetimes. To extract these lifetimes, fluorescence decays were fit to a two-component exponential decay that was re-convolved with the instrument response function:

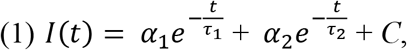

where I(t) = is the light intensity at time t following the laser pulse, τ_1_ and τ_2_ are the short (quenched) and long (unquenched) lifetimes of the fluorophore, and α_1_ and α_2_ are the fractional component of each lifetime. *C* is included to account for background light. For NAD(P)H, the short lifetime (τ_1_) corresponds to unbound NAD(P)H and the long lifetime (τ_2_) corresponds to protein-bound NAD(P)H (*29*). The opposite is true of FAD: the short and long lifetime correspond to bound FAD and unbound FAD, respectively (*24*). A mean lifetime at each pixel was also computed as the weighted average of the short and long lifetime:

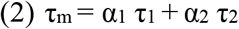

Following extraction of the fluorescence lifetimes, images were segmented to create single-cell masks using NAD(P)H intensity images. Segmentation was carried out in CellProfiler, resulting in masks of cells, cell nuclei, and cell cytoplasm. PerCP-conjugated CD69 fluorescence images were manually segmented by a trained observer. The observer was blinded to whether PerCP-CD69 images came from the activated or unstimulated condition. The resulting masks were used to identify activated and quiescent cells in each condition based on overlap between PerCP-CD69 masks and cell masks.

Fluorescent lifetime components for each cell were calculated in R. The values of NAD(P)H τ_m_, NAD(P)H τ_1_, NAD(P)H τ_2_, NAD(P)H α_1_, FAD τ_m_, FAD τ_1_, FAD τ_2_, and FAD α_1_ were calculated for each cell by averaging across all pixels in the cell cytoplasm. Cells with low photon counts (< 5000 photons), small masks that are unlikely cells (< 350 pixels or 75 μm^2^ whole cell area), and pixels with poor goodness-of-fit (χ^2^ > 1.3) were not included in this analysis. α_2_ was not computed, as the sum of α_1_ and α_2_ is equal to 1 (100%). An additional parameter, the optical redox ratio, was computed for each cell, defined here as the NAD(P)H intensity divided by the sum of the NAD(P)H and FAD intensities. This definition of the redox ratio is bound between 0 and 1. To account for variations in intensity from day-to-day equipment and setting changes, the redox ratio of each cell was normalized to the mean redox ratio of the control group within each experiment.

### 4.6 Measurement of cytokines and glucose/lactate levels in primary cell media

To validate the activation of lymphocytes in each condition, cytokine levels were measured in media samples collected from both the unstimulated and activated conditions during plating (24 hours post activation for NK cells, and 72 hours post activation for B cells). IFN-γ levels were measured in NK cell media samples using the human IFN-γ DuoSet ELISA kit (R&D Systems). IL-6 levels were measured in B cell media samples using the human IL-6 DuoSet ELISA kit (R&D Systems) (*53, 54*). The ELISA assay was carried out according to the provided protocol. Plates were incubated overnight with 2 μg/mL IFN-γ or IL-6 capture antibody. The plates were then washed and blocked with a 1% bovine serum albumin solution for 1 hour. Following washing, media samples and standards were incubated on the plates for 2 hours at room temperature, followed by a 2 hour incubation with 200 ng/mL IFN-γ or 50 ng/mL IL-6 detection antibody. Finally, the plates were incubated with streptavidin-conjugated horseradish peroxidase B, then an H2O2-tetramethylbenzidine substrate solution. The color reaction was stopped at 20 minutes with a 4M H2SO4 solution, and the plates were transferred to a plate reader, where they were read at 450 nm with wavelength correction at 570 nm. Standard curves were calculated from a serial dilution of the standards using a sigmoidal four parameter logistic model. The R^2^ of the standard curves for the IL-6 and IFN-γ ELISA experiments were 0.9993 and 0.9997, respectively.

To validate that the cells were upregulating aerobic glycolysis in the activated cell populations, commercial kits were used to measure glucose and lactate levels in media samples collected from both the unstimulated and activated conditions during plating (24 hours post activation for NK cells, and 72 hours post activation for B cells). A sample of the growth media used for the B cells and NK cells described in section 4.2 was also evaluated as a control. The glucose and lactate assays were carried out according to the respective protocols for the Glucose Colorimetric/Fluormetric Assay Kit (BioVision) or the Lactate Colorimetric/Fluormetric Assay Kit (BioVision). 0.5 μL of each sample was added to a 96-well plate were an additional 49.5 μL of assay buffer was added, yielding a 100x dilution of the original samples. 50 μL of reaction mix (2 μL probe, 2 μL enzyme mix, and 46 μL assay buffer) was then added to each well to yield a total volume of 100 μL per well. The 96-well trays were left to incubate for 30 minutes in a dark box at room temperature (glucose assay) or 37°C (lactate assay). The plates were then transferred to a plate reader where glucose or lactate levels were quantified by absorbance at OD 570. Standard curves were calculated from a serial dilution of the standards using an ordinary least squares regression model. The R^2^ of the standard curves for the glucose and lactate assays were 0.9973 and 0.9979, respectively.

### 4.7 Previous CD3^+^ T cell data

Previously published T cell data from Walsh. et. al. (2021) was used for the purposes of classifying lymphocyte subtypes in **Fig. 5**, **Supp. Figs. 5–8, 10**, and **Supp. Table 1** (*47*). T cells were isolated from human blood and either left quiescent or stimulated with CD2/3/28 for 48 hours for activation. Activation was confirmed with a CD69-PerCP label across three donors. This prior data was collected in the same manner on the same two-photon fluorescence lifetime imaging system as the current NK cell and B cell data.

### 4.8 Heatmap, UMAP, and Classification

Z-score heatmaps were constructed in R using the Complex Heatmap package (*55*). Clustering of groups or single cells was performed based on the OMI parameters and calculated using Ward’s method. Labels for activation, lymphocyte subtype, and donor were added afterwards and were not included in cluster analysis.

Uniform Manifold Approximation and Projection (UMAP) is a non-linear dimension reduction technique that can be used to visualize high-dimensional data. UMAP projections were made in Python using scikit-learn, UMAP, and Holoviews. Unless otherwise noted, each UMAP is a two-dimensional visualization of 9 variables (normalized optical redox ratio; NAD(P)H τ_m_, τ_1_, τ_2_, α_1_; FAD τ_m_, τ_1_, τ_2_, α_1_). The UMAP projection was computed using Euclidean distance. The nearest neighbors parameter was set to 15 and the minimum distance was set to 0.4 unless otherwise noted.

Random forest classification methods were trained in Python using scikit-learn to classify activation and/or lymphocyte subtype in the NK cell OMI parameters, the B cell OMI parameters, or combined OMI parameters from NK cells, B cells, and previously published T cell data (*47*). The classifier was trained on a random selection of 70% of the input data and tested on the remaining 30% for B cell or NK cell classifiers alone (*i.e*., Figs. 2, 4, and Supp. Figs. 2, 4). For the classifiers using a combined lymphocyte dataset of B, NK, and T cells (*i.e*., Fig. 5 and Supp. Figs. 5–8), the classifier was trained on a random selection of 50% of the input data and tested on the remaining 50%. Multiple metrics were used to evaluate the robustness of the classifier, including the receiver operating characteristic (ROC) curve, accuracy, precision, and recall. Classifiers were trained and tested on different random sets of the data to check for consistency in these metrics. Equal cost was given to a misclassified cell regardless of category (*i.e*., misclassification was not weighted by sample size).

Phasor-based classification was performed using the NAD(P)H and FAD phasor coordinates (G,S) at the laser repetition frequency (80 MHz) and its second harmonic (160 MHz) as features. The phasor coordinates were averaged pixel-wise over each cell mask using pixel intensities as weights to calculate cell-level phasor coordinates. Logistic regression classifiers with a logit link function and random forest classifiers with 100 decision trees were used, and the classifiers were trained on a random selection of 50% of the input data and tested on the remaining 50%. Again, equal cost was given to a misclassified cell regardless of category (*i.e*., misclassification was not weighted by sample size). Both the phasor and the fit analysis pipelines use the same raw FLIM data and cell masks to calculate cell-level phasor coordinates and fit parameters, respectively. However, the exclusion criteria for the two pipelines are not the same, which results in different final number of cells included in the phasor-based and fit-based classifiers. For example, the phasor pipeline removes low-count (with fewer than 5000 photons) or small (with fewer than 50 pixels) cells, while the fit analysis also removes cells based on the goodness of the bi-exponential fit (χ^2^ > 1.3).

### 4.9 Statistical analysis

Statistical analysis was performed using the statannotations package v0.5.0 in Python.

Differences between groups were tested using Kruskal-Wallis with post-hoc comparisons test for multiple group comparisons, or a two-tailed unpaired T-test for comparisons of pairs of data.

## Acknowledgements

The authors would like to thank Dr. Jose Maria Ayuso Dominguez for advice and assistance with NK cell culture, Steve John (Jing Fan Lab) for assistance with the ELISA assays, Matthew Stefely for assistance with figure illustration, and Dr. Jens Eickhoff for statistical guidance.

## Funding

This research was supported by the

National Institutes of Health grant U24AI152177 (MCS)

National Institutes of Health grant R01CA211082 (MCS)

National Institutes of Health grant R01 CA205101 (MCS)

Morgridge Institute for Research (MCS)

Carol Skornicka Chair of Biomedical Imaging (MCS)

Retina Research Foundation Daniel M. Albert Chair (MCS)

National Science Foundation Center for Cell Manufacturing Technologies grant EEC-1648035 (MCS, CMC)

National Institutes of Health grant CA215461 (CMC)

St. Baldrick’s EPICC Team Translational Research Grant (MHF, CMC)

The MACC Fund (AS, CMC).

The University of Wisconsin (UW) Carbone Cancer Center Flow Cytometry Laboratory is supported by NIH/NCI P30 CA014520.

The contents of this article do not necessarily reflect the views or policies of the Department of Health and Human Services, nor does mention of trade names, commercial products, or organizations imply endorsement by the US Government.

## Author Contributions

AJW and MCS developed the central hypotheses and designed the study. AJW, KT, and IJ performed the B cell OMI experiments and analyzed the data. RLS, JR, MHF, AS, and EMM performed the NK cell OMI experiments and analyzed the data. RLS performed the ELISA experiments. JR performed the glucose and lactate analysis. RLS, KS, ECG, and JR put together figures. PR and ECG provided code for the UMAPs and random forest classifiers, which was applied by PR, ECG, KS, and RLS. KS performed the phasor-based classification. MCS and CMC provided project guidance and oversight. RLS wrote the first draft of the manuscript. All authors provided input into the final version of the manuscript.

## Data and materials availability

All data and code used in the analyses is available for purposes of reproducing or extending the analyses through a GitHub repository (https://github.com/skalalab/schmitz_r-lymphocyte_activation).

---

**Supplemental Figure 1.**
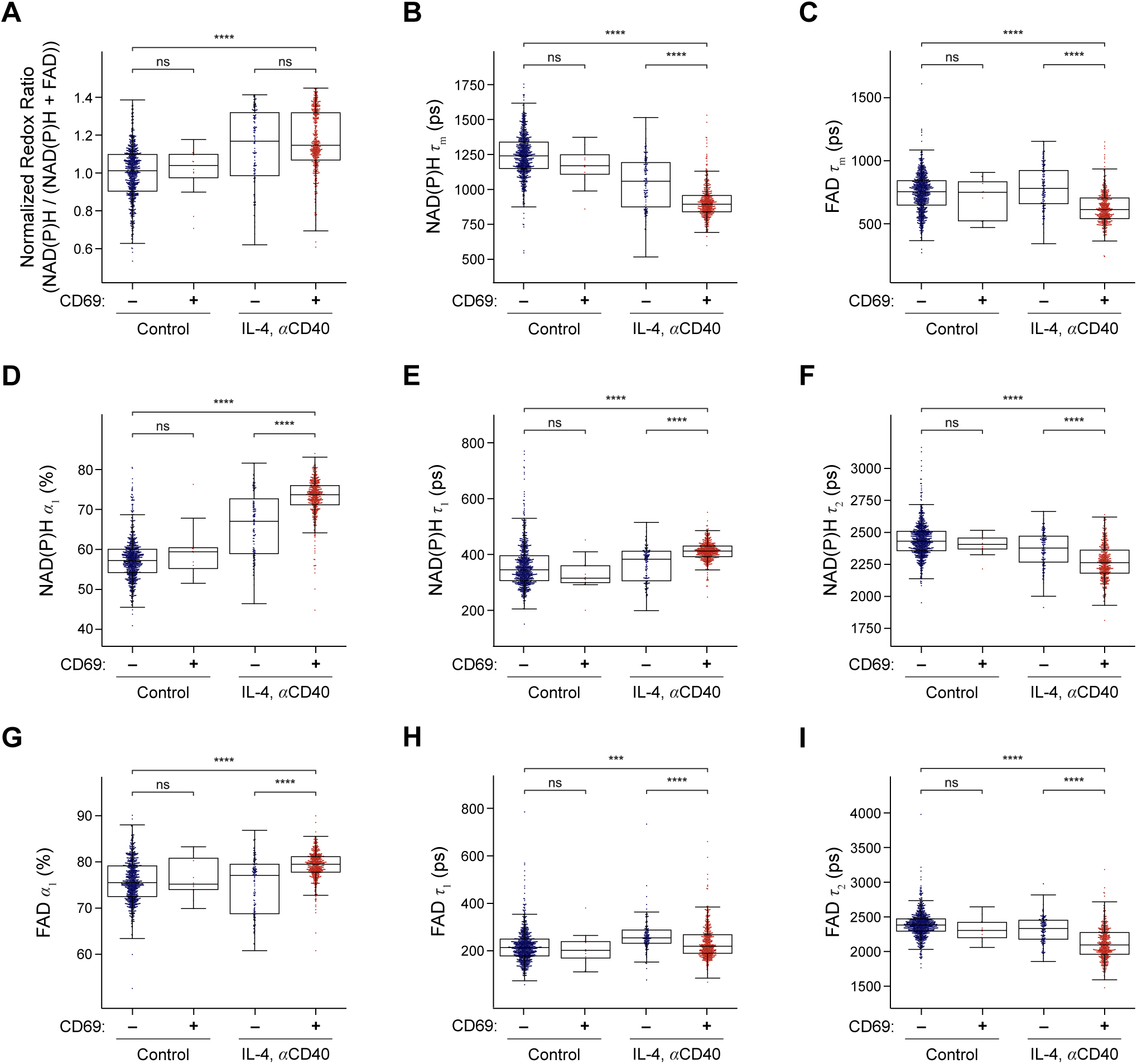
OMI of CD69+ and CD69-B cells in both control and anti-CD40 + IL-4 activated culture. Both CD69+ and CD69-B cells in both conditions (control and IL-4 + anti-CD40 activated) for each OMI parameter: (A) optical redox ratio (NAD(P)H intensity divided by the sum of NAD(P)H + FAD intensities), (B) NAD(P)H mean lifetime τ_m_, (C) FAD mean lifetime τ_m_, (D) unbound NAD(P)H fraction α_1_, (E) unbound NAD(P)H lifetime τ_1_, (F) protein-bound NAD(P)H lifetime τ_2_, (G) protein-bound FAD fraction α_1_, (H) protein-bound FAD lifetime τ_1_, (I) unbound FAD lifetime τ_2_. Plots display single cell values (dots) overlaid on box-and whisker plots displaying the median, interquartile range (IQR), and minimum/maximum value. n = 1352 (461 cells in the activated CD69+ condition, 130 cells in the activated CD69-condition, 12 cells in the control CD69+ condition, 749 cells in the control CD69-condition). *** p < 0.005, **** p < 0.0001, Kruskal-Wallis with post-hoc comparisons. ns = not significant.

**Supplemental Figure 2.**
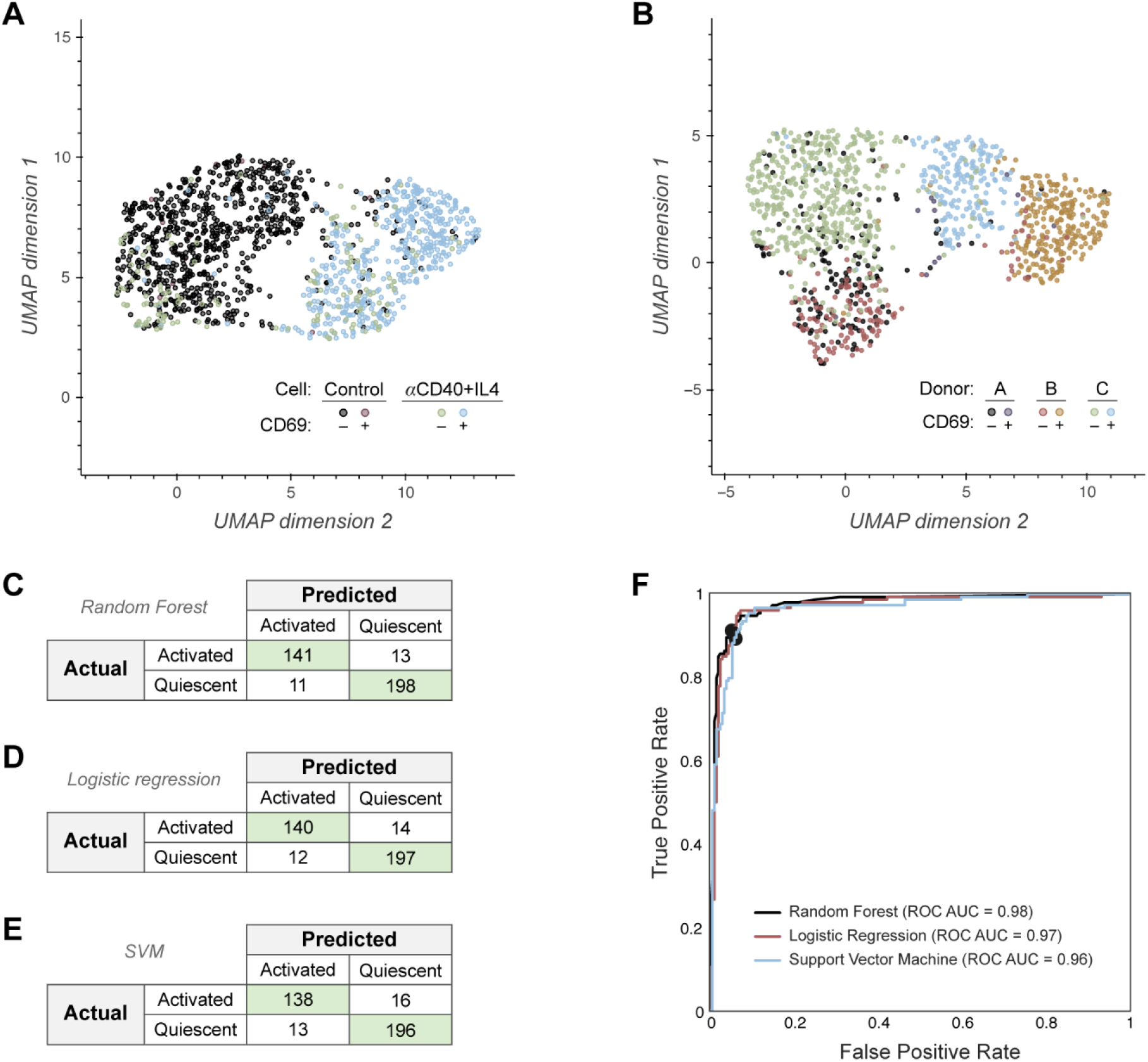
Additional UMAPs and classifier performance for single B cell OMI. (A) UMAP of B cells with labels for both CD69+ and CD69-cells in the control and activated (anti-CD40+IL4) groups. (B) UMAP of B-cells color-coded by donor (A, B, C) and activation status (CD69+, CD69-). (C) Confusion matrix of the 9 OMI parameter random forest classifier shows performance for classification of CD69+ activated and CD69-control B cells. (D) Confusion matrix of a logistic regression classifier trained on 9 OMI parameters to classify B cells as CD69+ activated or CD69-control. (E) Confusion matrix of a support vector machine (SVM) classifier trained on 9 OMI parameters to classify B cells as CD69+ activated or CD69-control. (F) ROC curves for random forest, logistic regression, and SVM classifiers trained on 9 OMI parameters, with operating points indicated. In (A), n = 1352 (461 cells in the activated CD69+ condition, 130 cells in the activated CD69-condition, 12 cells in the control CD69+ condition, 749 cells in the control CD69-condition). In (B) – (F), n = 1210 cells (461 cells in the activated CD69+ condition, 749 cells in the control CD69-condition) with a 70/30 split for training and test sets.

**Supplemental Figure 3.**
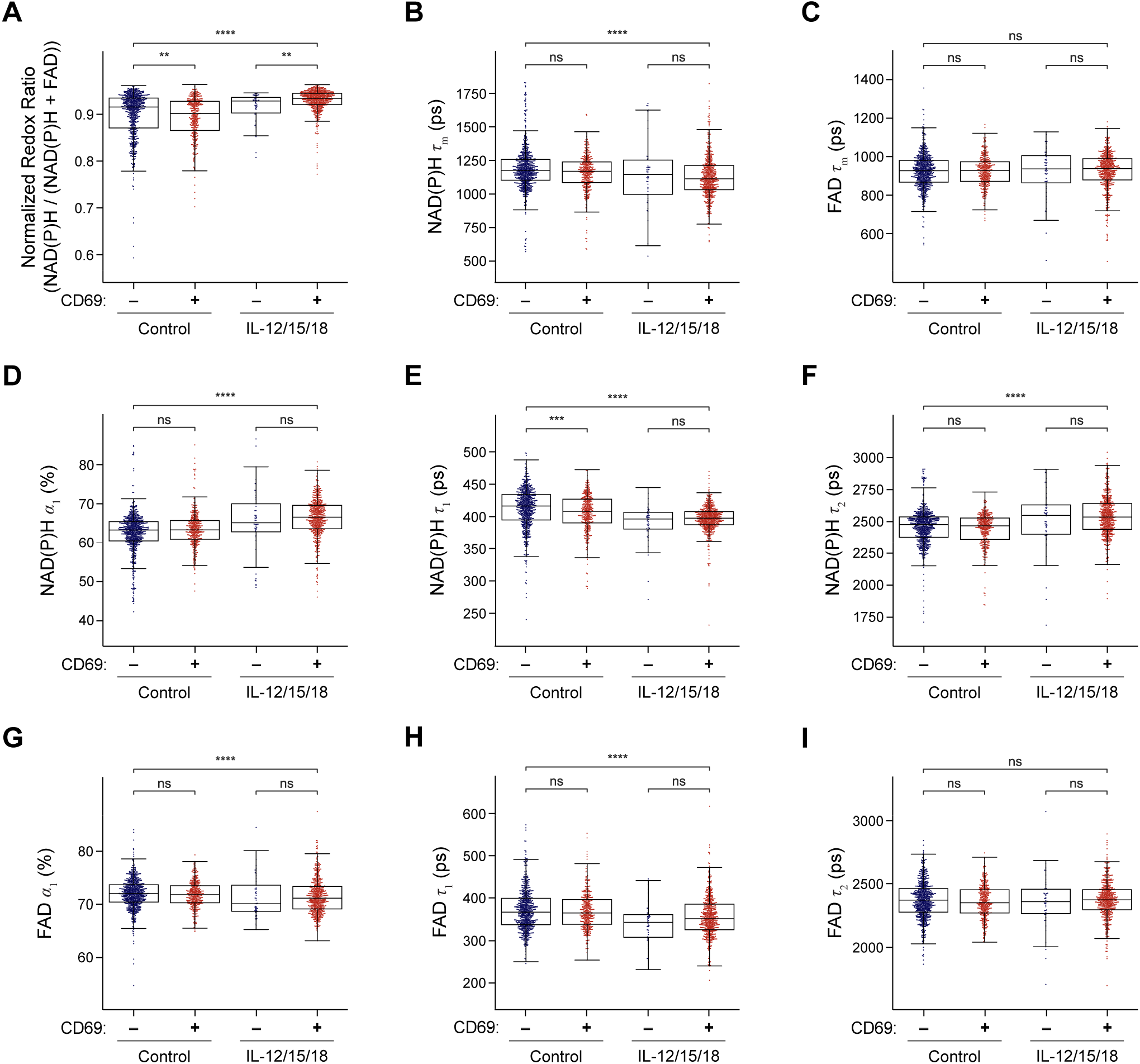
OMI of CD69+ and CD69-NK cells in both unstimulated and IL-12 + IL-15 + IL-18 activated culture conditions. Both CD69+ and CD69-NK cells in both conditions (control and IL-12 + IL-15 + IL-18 activated) for each OMI parameter: (A) optical redox ratio, (B) NAD(P)H mean lifetime τ_m_, (C) FAD mean lifetime τ_m_, (D) unbound NAD(P)H fraction α_1_, (E) unbound NAD(P)H lifetime τ_1_, (F) protein-bound NAD(P)H lifetime τ_2_, (G) protein-bound FAD fraction α_1_, (H) protein-bound FAD lifetime τ_1_, (I) unbound FAD lifetime τ_2_. Plots display single cell values (dots) overlaid on box-and whisker plots displaying the median, interquartile range (IQR), and minimum/maximum value. n = 1642 cells (554 activated CD69+ cells, 372 activated CD69-cells, 49 control CD69+ cells, 667 control CD69-cells). ** p < 0.01, *** p < 0.001, **** p < 0.0001, Kruskal-Wallis with post-hoc comparisons. ns = not significant

**Supplemental Figure 4.**
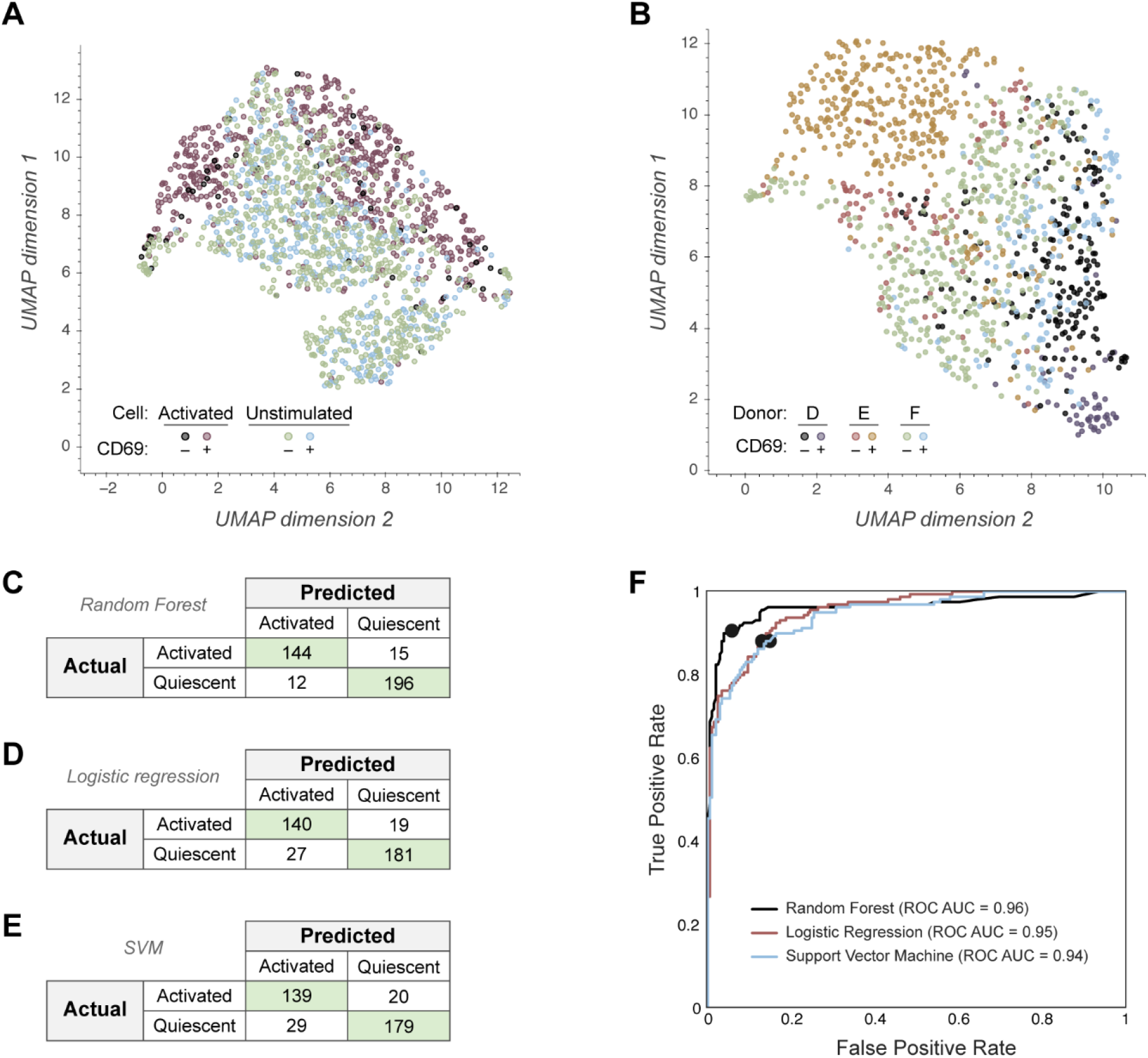
Additional UMAPs and classifier performance for single NK cell OMI. (A) UMAP of NK cells with labels for both CD69+ and CD69-cells in the control and activated (IL-12 + IL-15 + IL-18) groups. (B) UMAP of NK cells color-coded by donor (D, E, F) and activation status (CD69+, CD69-). (C) Confusion matrix of the 9 OMI parameter random forest classifier trained to classify NK cells as CD69+ activated or CD69-control cells. (D) Confusion matrix of a logistic regression classifier trained on 9 OMI parameters to classify NK cells as CD69+ activated or CD69-control. (E) Confusion matrix of SVM classifier trained on 9 OMI parameters to classify NK cells as CD69+ activated or CD69-control. (F) ROC curves for the random forest, logistic regression, and SVM classifiers trained on 9 OMI parameters, with operating points indicated. In (A), n = 1642 cells (554 activated CD69+ cells, 372 activated CD69-cells, 49 control CD69+ cells, 667 control CD69-cells). In (B) – (F), n = 1221 cells (554 cells in the activated CD69+ condition, 667 cells in the control CD69-condition) with a 70/30 split for training and test sets.

**Supplemental Figure 5.**
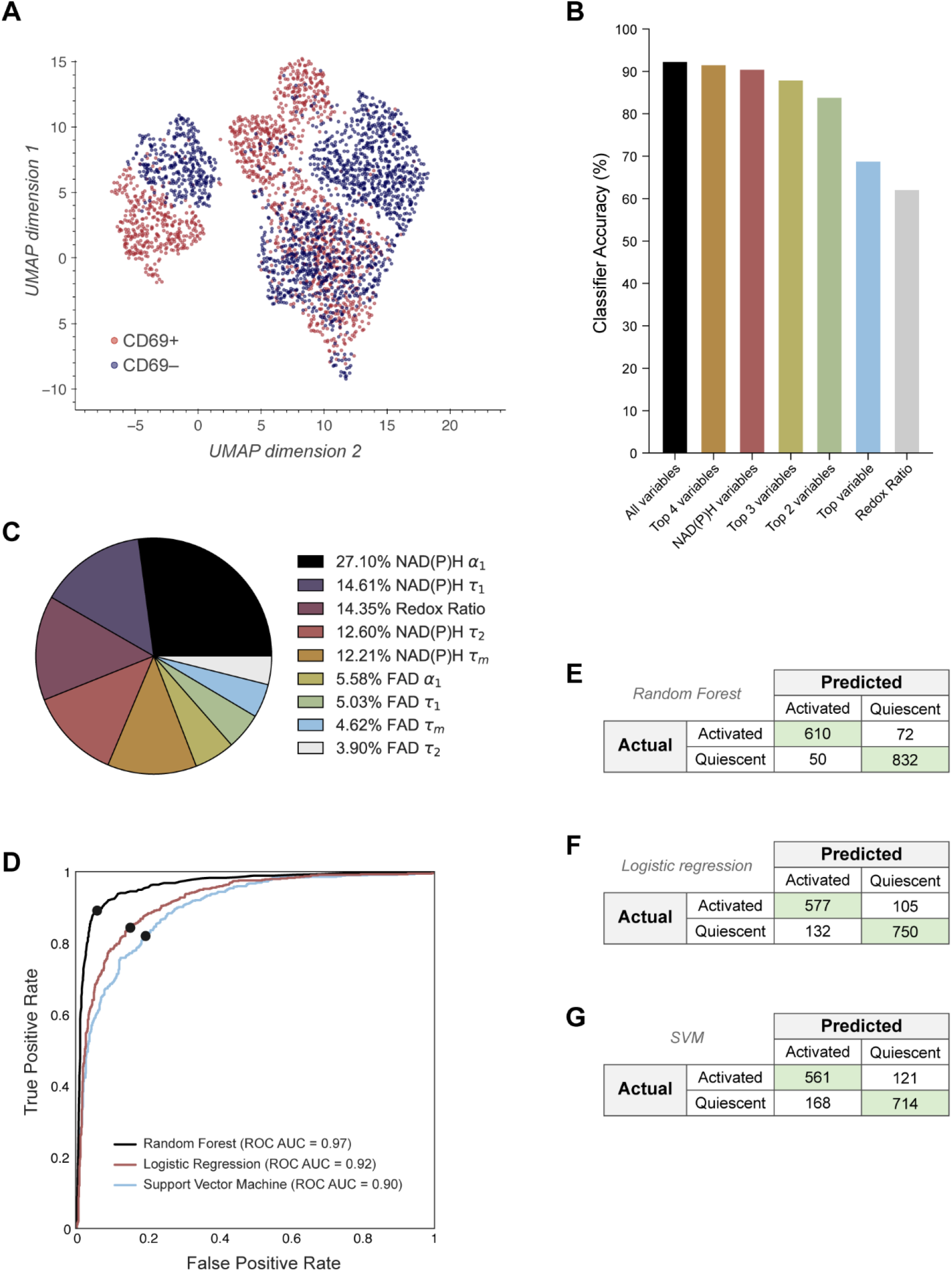
Additional UMAPs and classifier performance for activation of lymphocytes (T cells, B cells and NK cells). (A) UMAP of single-cell OMI data from Fig. 5C containing all T, B, and NK cells color-coded by activation status. (B) Bar graph of % accuracy for a random forest classifier trained to distinguish CD69+ from CD69-cells across the combined dataset of all lymphocyte subtypes. (C) Pie chart displaying the weights of OMI variables included in the random forest classifier using all 9 OMI features in Fig. 5D. (D) ROC curves of random forest, logistic regression, and support vector matrix (SVM) classifiers using all 9 OMI variables to distinguish CD69+ from CD69-cells across all lymphocyte subtypes, with operating points indicated. (E) Confusion matrix for random forest classifier using all 9 OMI variables to classify cells as activated (CD69+) or quiescent (CD69-) in Fig. 5D. (F) Confusion matrix for logistic regression classifier. (G) Confusion matrix for SVM classifier. n = 3127 cells (1747 CD69-cells, 1380 CD69+ cells) with a 50/50 split for training and test sets. T cell data taken from previously published dataset (*47*).

**Supplemental Figure 6.**
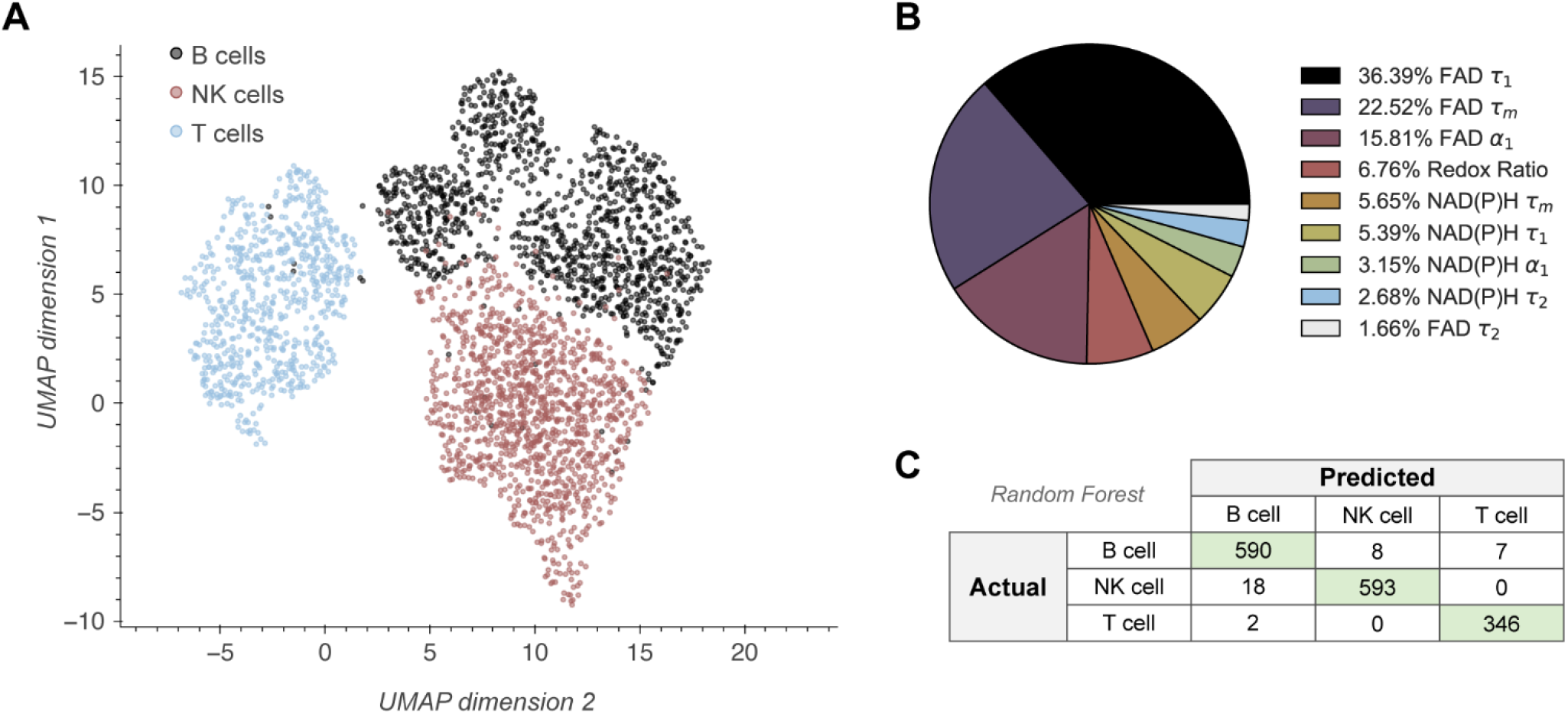
Additional UMAPs and classifier performance for lymphocyte subtype (T cells, B cells and NK cells). (A) UMAP of lymphocytes from Fig. 5C color-coded by lymphocyte subtype. (B) Variable weights of 9 OMI parameters used for one-vs.-one random forest classification by lymphocyte subtype in Fig. 5E. (C) Confusion matrix for 9 OMI parameter random forest classifier in Fig. 5E. n = 3127 cells (1210 B cells, 1221 NK cells, 696 T cells) with a 50/50 split for training and test sets. T cell data taken from previously published dataset (*47*).

**Supplemental Figure 7.**
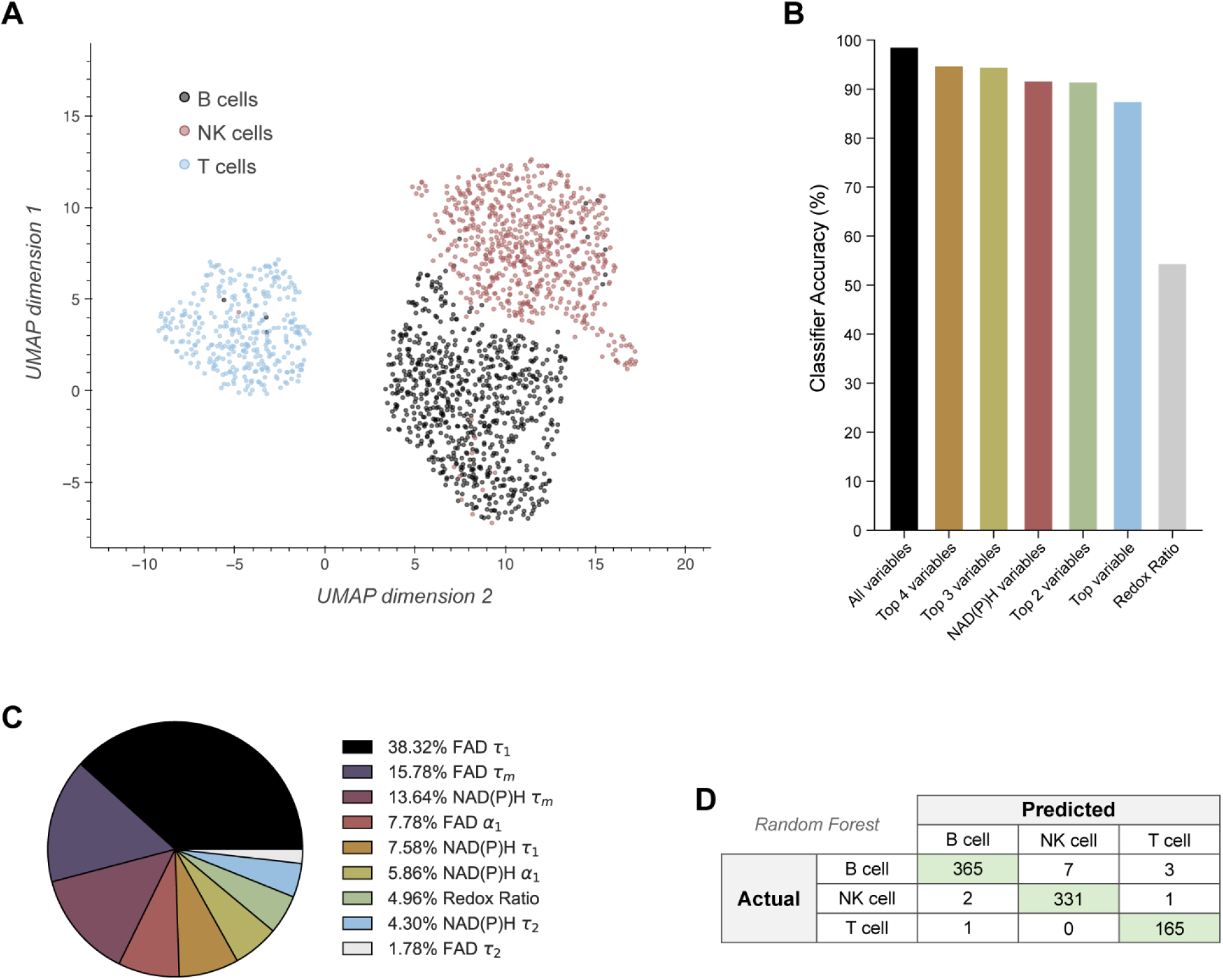
UMAP and classifier performance for lymphocyte subtype classifier based on quiescent cells only (T cells, B cells and NK cells). (A) UMAP of lymphocytes from quiescent (CD69-control) NK, B, and T cells color-coded by lymphocyte subtype. (B) Bar graph displaying accuracy of random forest classifiers trained to separate lymphocytes based on lymphocyte subtype (one vs. one approach, quiescent cells only). (C) Feature weights of 9 OMI parameters used for one-vs.-one random forest classification by lymphocyte subtype in (B). (D) Confusion matrix for 9 OMI parameter random forest classifier in (B). n = 1747 cells (749 B cells, 667 NK cells, 331 T cells) with a 50/50 split for training and test sets. T cell data taken from previously published dataset (*47*).

**Supplemental Figure 8.**
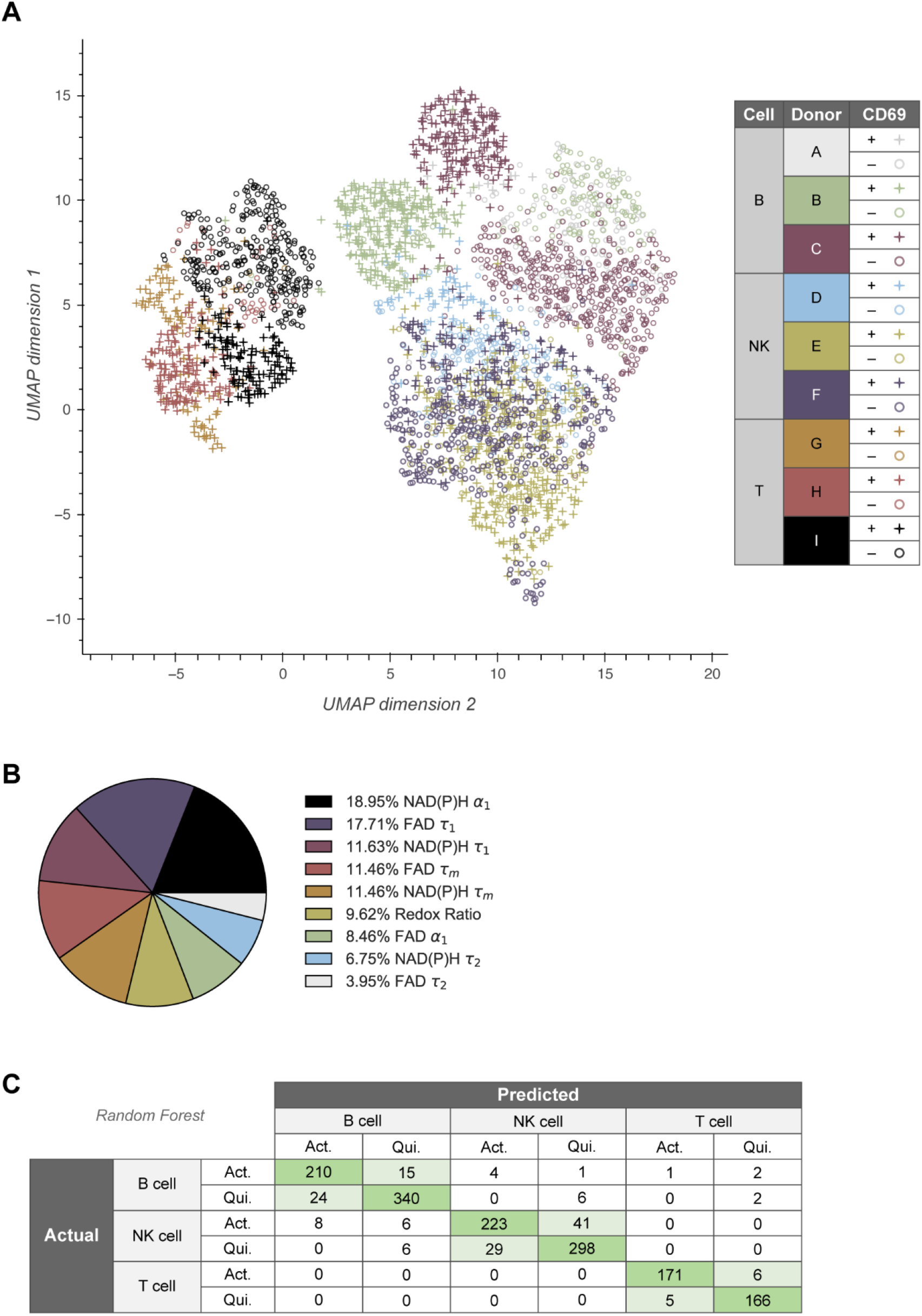
Additional UMAPs and classifier performance for both lymphocyte subtype (T cells, B cells and NK cells) and activation. (A) UMAP of lymphocytes from Fig. 5C color-coded by lymphocyte subtype, activation status, and donor. (B) Feature weights of 9 OMI parameters used for one-vs.-one random forest classification by lymphocyte subtype and activation status in Fig. 5F (C) Confusion matrix for 9 OMI parameter random forest classifier in Fig. 5F. n = 3127 cells (749 CD69-control B cells, 461 CD69+ activated B cells, 667 CD69-control NK cells, 554 CD69+ activated NK cells, 331 CD69-control T cells, 365 CD69+ activated T cells) with a 50/50 split for training and test sets. T cell data taken from previously published dataset (*47*).

**Supplemental Figure 9.**
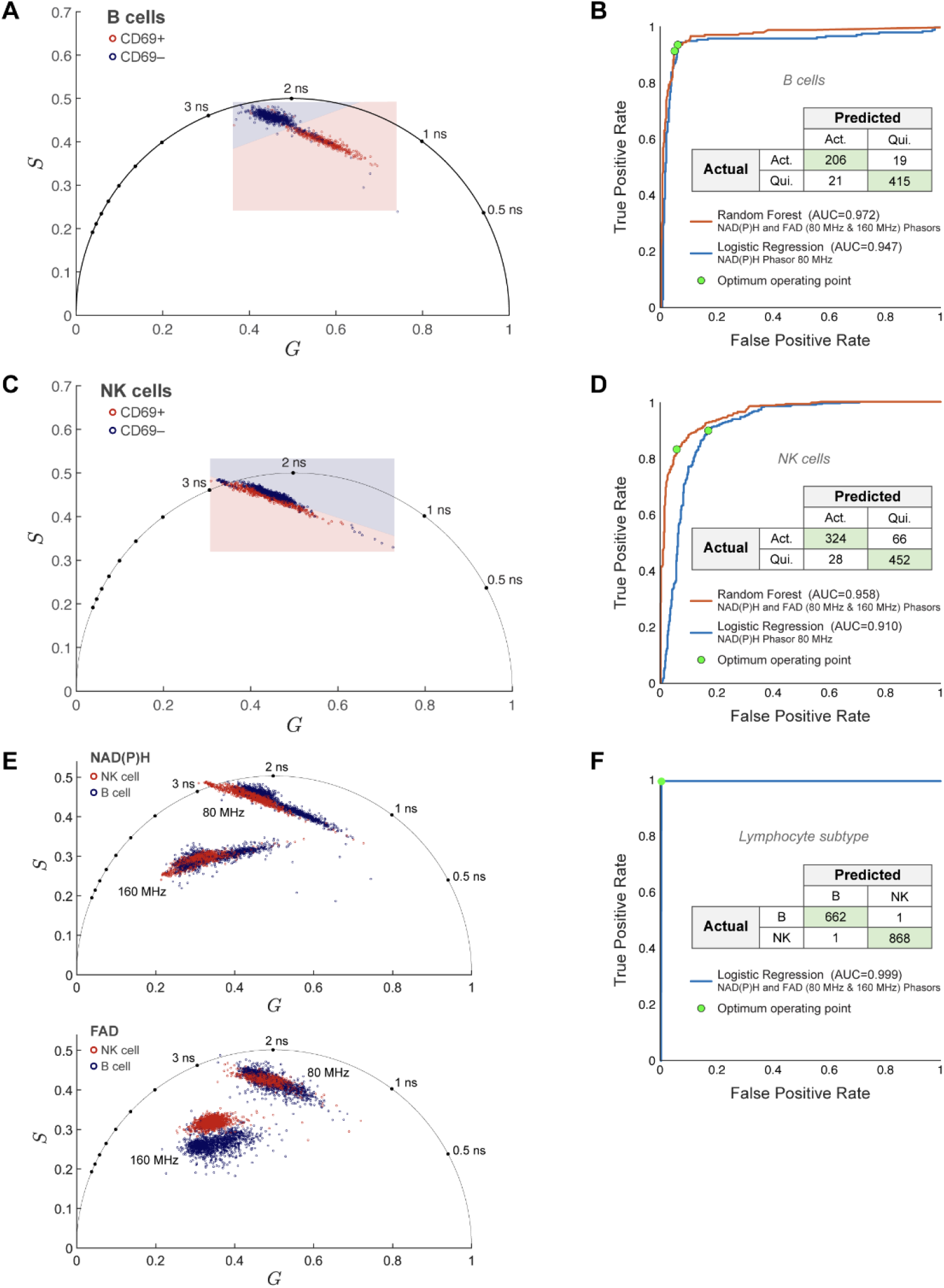
Phasor-based classification of NK cell and B cell activation and lymphocyte subtype. (A) NAD(P)H phasor plot of B cells from Fig. 1 (Red = activated CD69+ B cells, blue = quiescent CD69-B cells). Shaded areas show decision boundaries for logistic regression classification of B cell activation based on NAD(P)H phasor. (B) ROC curves and confusion matrix for random forest classification of B cell activation. The NAD(P)H and FAD phasors at both the laser repetition frequency (80MHz) and its second harmonic (160MHz) predicted B cell activation with a classification accuracy of 93.9%, n = 1323 B cells (451 cells in the activated CD69+ condition, 872 cells in the control CD69-condition) with a 50/50 split for training and test sets. (C) NAD(P)H phasor plot of NK cells from Fig. 3 (Red = activated CD69+ NK cells, blue = quiescent CD69-NK cells). Shaded areas show decision boundaries for logistic regression classification of NK cell activation based on NAD(P)H phasor. (D) ROC curves and confusion matrix for random forest classification of NK cell activation. The NAD(P)H and FAD phasors at both the laser repetition frequency (80MHz) and its second harmonic (160MHz) predicted NK cell activation with a classification accuracy of 89.2%, n = 1742 cells (781 cells in the activated CD69+ condition, 961 cells in the control CD69-condition) with a 50/50 split for training and test sets. (E) Phasor plots of NAD(P)H (top) and FAD (bottom) of B cells and NK cells at both the laser repetition rate (80 MHz) and its second harmonic (160 MHz). (F) ROC curve and confusion matrix for logistic regression classification of B vs NK. Using both the NAD(P)H and FAD phasors at 80MHz and 160MHz, the logistic regression model could classify a cell as B or NK with a classification accuracy of 99.9%, n = 3065 cells (1323 B cells, 1742 NK cells) with a 50/50 split for training and test sets. Due to separate processing pipelines, the total number of cells in phasor analysis differs from that in the fit analysis. Both analysis pipelines use the same raw data.

**Supplemental Figure 10.**
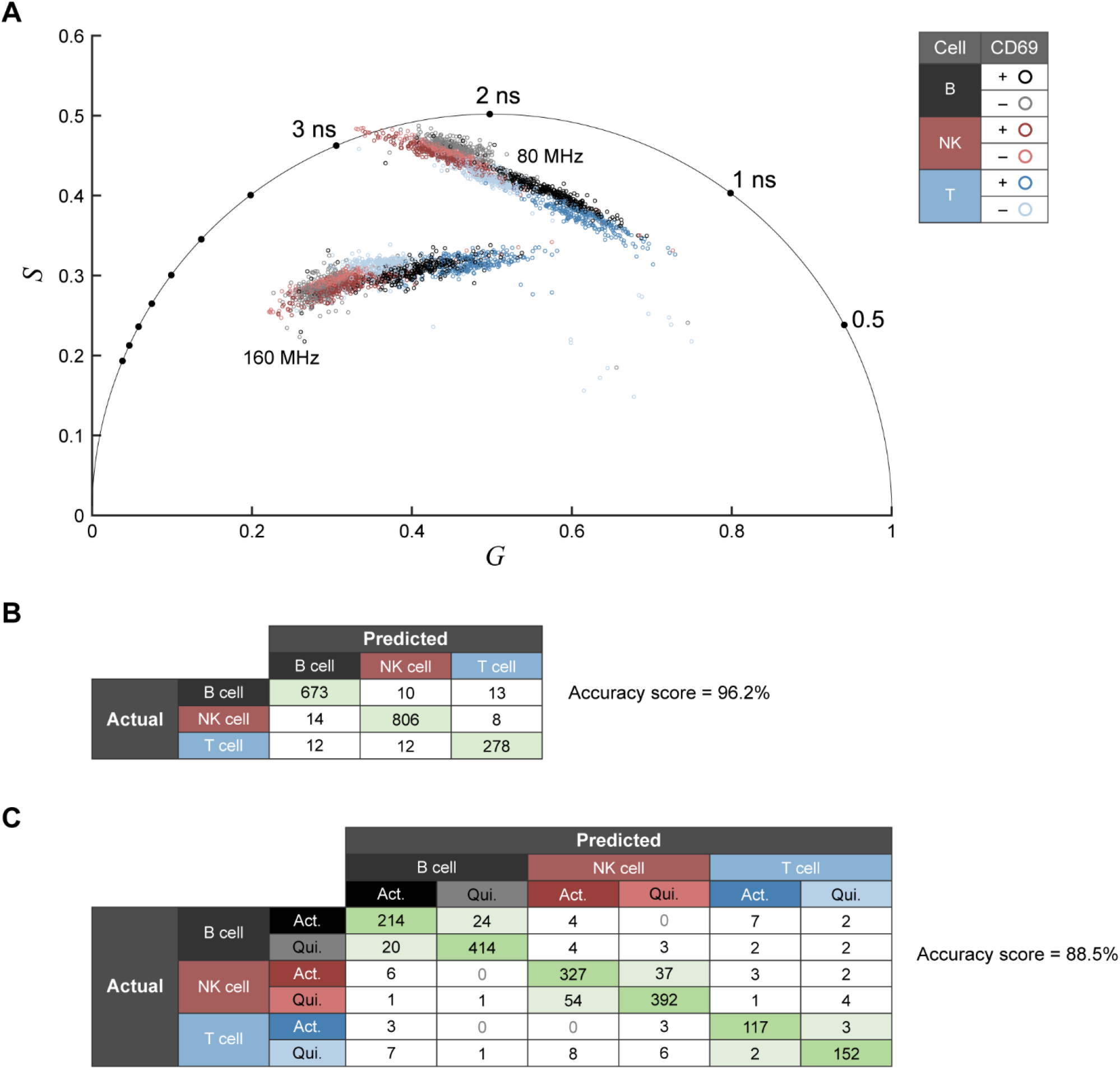
Phasor-based classification of NK cell, B cell, T cell lymphocyte subtype and activation. (A) NAD(P)H phasor plot of B cells, NK cells, and T cells. (B) Confusion matrix for random forest classification of lymphocyte subtype. The classifier was trained on NAD(P)H phasors (at the laser repetition frequency 80MHz and its second harmonic 160 MHz) and achieves a classification accuracy of 96.2%. (C) Confusion matrix for random forest classification of lymphocyte subtype and activation trained on NAD(P)H phasors achieves a classification accuracy of 88.5%. The classifier predicted lymphocyte subtype and activation with a total n=3653 lymphocytes including n = 1323 B cells (451 B cells in the activated CD69+ condition, 872 B cells in the control CD69-condition); n = 1742 NK cells (781 cells in the activated CD69+ condition, 961 cells in the control CD69-condition); n = 588 T cells (263 cells in the activated condition, 325 cells in the control condition); a 50/50 data split for training and test sets was used. Due to separate processing pipelines and exclusion criteria, the total number of cells in phasor analysis differs from that in the fit analysis. Both analysis pipelines use the same raw data. T cell data taken from previously published dataset (*47*).

**Supplemental Table 1:** Accuracies for random forest classifiers given in main and supplemental figures. Accuracies are given out of a maximum of 100%, where accuracy = total number of correct classifications / total number of all classifications * 100. “Top” 1, 2, 3, and 4 variables classifiers refer to the largest weighted variables in the “all variable” classifier, found in the corresponding figure (or Supp. Fig. 6B and 8B for Main Fig. 5E and 5F, respectively). In Main Fig. 4D, the top variable is the optical redox ratio (RR). NAD(P)H column refers to classifiers that used all NAD(P)H lifetime variables (NAD(P)H τ_m_, τ_1_, τ_2_, α_1_ as inputs. ALL, all lymphocytes including T, B, NK cells; ALL-qui, quiescent only lymphocytes including T, B, NK cells. T cell data taken from previously published dataset (*47*).

| Figure | Cell Type | Classification | Variables |  |  |  |  |  |  |  |
| --- | --- | --- | --- | --- | --- | --- | --- | --- | --- | --- |
| | | | All | Top 1 | Top 2 | Top 3 | Top 4 | NAD(P)H | RR | NAD(P)H $\alpha_1$ |
| Main 2D | B | activation | 93.39% | 88.98% | 92.01% | 93.11% | 93.39% | 92.56% | 63.36% |  |
| Main 4D | NK | activation | 92.64% | 61.58% | 81.20% | 89.37% | 92.37% | 91.55% | *top variable* |  |
| Main 5E | ALL | lymphocyte subtype | 97.76% | 85.74% | 90.66% | 92.84% | 96.42% | 89.90% | 56.33% | 64.00% |
| Main 5F | ALL | lymphocyte subtype and activation | 90.03% | 50.45% | 74.36% | 81.27% | 83.18% | 83.31% | 37.02% |  |
| Supp. 5B | ALL | activation | 92.20% | 68.73% | 83.76% | 87.85% | 91.43% | 90.35% | 62.02% |  |
| Supp. 7B | ALL-qui. | lymphocyte subtype | 98.40% | 87.31% | 91.31% | 94.40% | 94.63% | 91.54% | 54.29% |  |

